# A Unified Atlas of T cell Glycophysiology

**DOI:** 10.1101/2024.08.24.609521

**Authors:** Fauzia N. Izzati, Hani Choksi, Paolo Giuliana, Diala Abd-Rabbo, Heidi Elsaesser, Aled Blundell, Vanessa Affe, Vinicius Kannen, Zeinab Jame-Chenarboo, Edward Schmidt, Meggie Kuypers, David B. Avila, Eric S. Y. Chiu, Dzhangar Badmaev, Haissi Cui, Jason Matthews, Thierry Mallevaey, Matthew S. Macauley, David G. Brooks, Landon J. Edgar

**Author notes:** These authors contributed equally to this work (Fauzia N. Izzati, Hani Choksi).

## Abstract

Glycans are emerging as important regulators of T cell function but remain poorly characterized across the functionally distinct populations that exist *in vivo*. Here, we couple single-cell analysis technologies with soluble lectins and chemical probes to interrogate glycosylation patterns on major T cell populations across multiple mouse and human tissues. Our analysis focused on terminal glycan epitopes with immunomodulatory functions, including sialoglycan ligands for Siglecs. We demonstrate that glycosylation patterns are diverse across the resting murine T cell repertoire and dynamically remodelled in response to antigen-specific stimulation. Surprisingly, we find that human T cell populations do not share the same glycoprofiles or glycan remodelling dynamics as their murine counterparts. We show that these differences can be explained by divergent regulation of glycan biosynthesis pathways between the species. These results highlight fundamental glycophysiological differences between mouse and human T cells and reveal features that are critical to consider for glycan-targeted therapies.

## INTRODUCTION

T cells are central mediators of adaptive immunity and provide durable protection against pathogens and cancer.^1^ Activation of naïve T cells (T_N_) through T cell receptor (TCR) clustering, relay of co-stimulatory signals, and cytokine exposure induces dramatic remodelling of the T cell surface proteome.^2,3^ This results in unique protein expression profiles for differentiated T cell populations such as effector/memory (T_EM_) and central memory (T_CM_) cells.^4^ While protein-centered features of T cell activation have been well characterized for decades, they represent only a portion of the molecular landscape of the T cell surface. Like all cells in every organism, T cells are functionalized with a dense matrix of structurally diverse carbohydrate-protein/-lipid conjugates called glycans, which collectively form the glycocalyx.^5^ While glycans are now recognized as important regulators of T cell development^6–11^, the specific glycan repertoire and functional importance of these molecules to peripheral T cell immunophysiology remains poorly characterized.

Glycans on the surface of naïve murine T cells are remodelled following TCR stimulation and many of these changes^12–17^ involve glycans that terminate in monosaccharides belonging to the sialic acid family of sugars – collectively called sialoglycans (SGs). Sialic acids include (but are not limited to) *N*-glycolylneuraminic acid (Neu5Gc, not made in humans but common in mouse) and *N*-acetylneuraminic acid (Neu5Ac, found in both humans and mice). Neu5Gc or Neu5Ac can be installed on an underlying glycan core through multiple chemical linkages – α2-3- or α2-6-linked to galactose, α2-6-linked to *N*-acetylgalactosamine, or α2-8- to another sialic acid.^5,18^ In all cases, the privileged terminal position of sialic acid makes SGs important mediators of diverse biology that occurs at the cell surface, including cell:cell adhesion^19^ and immunoregulatory receptor recruitment at an immunological synapse – for example, engagement of immunoinhibitory sialic acid binding immunoglobulin-like lectins (Siglecs).^20,21^ Levels of SGs decrease following *ex vivo* stimulation of murine T cells^13^ – a process that enriches the T cell surface in galactose-terminating asialoglycans (ASG) and promotes apoptosis.^13,22–25^ Importantly, a growing body of work is now demonstrating that removal of sialic acids from glycan cores using recombinant neuraminidase enzymes promotes clonal expansion of T_N_s and augments effector functions of T_EM_s.^23,26–31^ Further, recent work has shown that disallowing SG biosynthesis through genetic knockout of the enzyme CMP-sialic acid synthetase (CMAS) enriches the murine T cell repertoire in highly-activated T_EM_ and T_CM_ populations.^32^ Discovery of these immunoregulatory features of the T cell glycocalyx has catalyzed interest in glycan editing technologies to control T cell immunity for therapeutic benefit.^31,33,34^ To this end, neuraminidase treatment of T cells and/or target cancer cells has been shown to potentiate anti-tumour activity in both murine and human systems.^31,34–36^ These results highlight the functional importance of terminal glycan epitopes on T cells and provide motivation to further characterize their diversity and dynamics during an immune response.

While glycan editing provides new dimensions of control over T cell-mediated immunity, we do not know which terminal glycan epitopes are presented on the discrete T cell populations that exist *in vivo* – for example, T_N_, T_EM_, T_CM_, or exhausted cells. Such information is needed to fully understand which cell types can be targeted by glycan editing technologies. Moreover, due to the lack of data defining glycosylation in resting or activated human T cells, it is unclear if murine T cell glycosylation patterns are mirrored by their human counterparts. Since targeting glycosylation is emerging as a strategy for human therapies, this information will be critical to guide translational efforts to edit T cell glycans for therapeutic benefit, especially when choosing model systems relevant to human immunophysiology. Here, we address these open questions using spectral flow cytometry and transcriptomic workflows to curate an atlas of terminal glycan epitopes on T cells from both mice and humans. Using soluble lectin probes, including recombinant Siglecs, we demonstrate that glycosylation is linked to phenotype and function across murine T cell differentiation and functional states. Importantly, we found that remodelling of SGs and ASGs on stimulated T cells was predominantly driven by changes in biosynthesis rather than catabolism of existing glycan structures. Unexpectedly, we observed that glycan remodelling events on stimulated human T cells were distinct from those that occurred on murine T cell counterparts, especially in the trajectories of α2-6-linked SG epitopes and presentation of specific glycan ligands for multiple Siglecs. These contrasting results could not be explained by differences in age, antigen experience, or tissues sampled and thus represent fundamental differences in T cell glycophysiology between the species. Taken together, our results furnish the first cross-species resource that describes T cell glycosylation in health and disease. These data will be broadly useful for improving technologies designed to target glycans on human T cells for therapeutic benefit.

## RESULTS

### The murine T cell glycocalyx is heterogeneous across functionally important populations

Our first goal was to assign glycosylation profiles to the individual T cells populations that exist in C57BL/6J specific pathogen-free (SPF) mice since they have been used as model systems for previous work in this space^9,13,31^. We used fluorescent plant-derived lectins with known terminal glycan-binding specificities^37^ in a spectral flow cytometry workflow which enabled us to catalogue α2-3- (using *Maackia amurensis* lectin, MAA-II) and α2-6- (using *Sambucus nigra* lectin, SNA) SGs in addition to galactose-terminating ASGs (*N*-acetyllactosamine (LacNAc) using *Erythrina crista-galli* lectin, ECL, and O-linked T antigen (Galβ1-3GalNAcα1-Ser/Thr) using peanut agglutinin, PNA) (**Figures 1A** and **1B**). These lectins are useful for monitoring the key SG and ASG epitopes that are remodelled during murine T cell activation and modified by neuraminidase technologies. We observed similar glycosylation patterns on T_N_ (CD44^Low^ CD62L^High^), T_CM_ (CD44^High^ CD62L^Low^), and pre-effector-like^38^ T cells (T_P4_, CD44^Low^ CD62L^Low^) in both CD4^+^ and CD8^+^ compartments; however, CD4^+^ and CD8^+^ T_EM_ (CD44^High^ CD62L^Low^) cells diverged into two populations based on α2-6-SG presentation (**Figures 1B, 1C** and **S1A**–**C**). T_EM_s also displayed higher levels of ASG structures, consistent with decreased termination of glycans in sialic acid residues. These results were recapitulated in T cells from murine peripheral blood samples (**Figure S1D**), indicating conserved glycosylation patterns across tissues.

**Figure 1:**
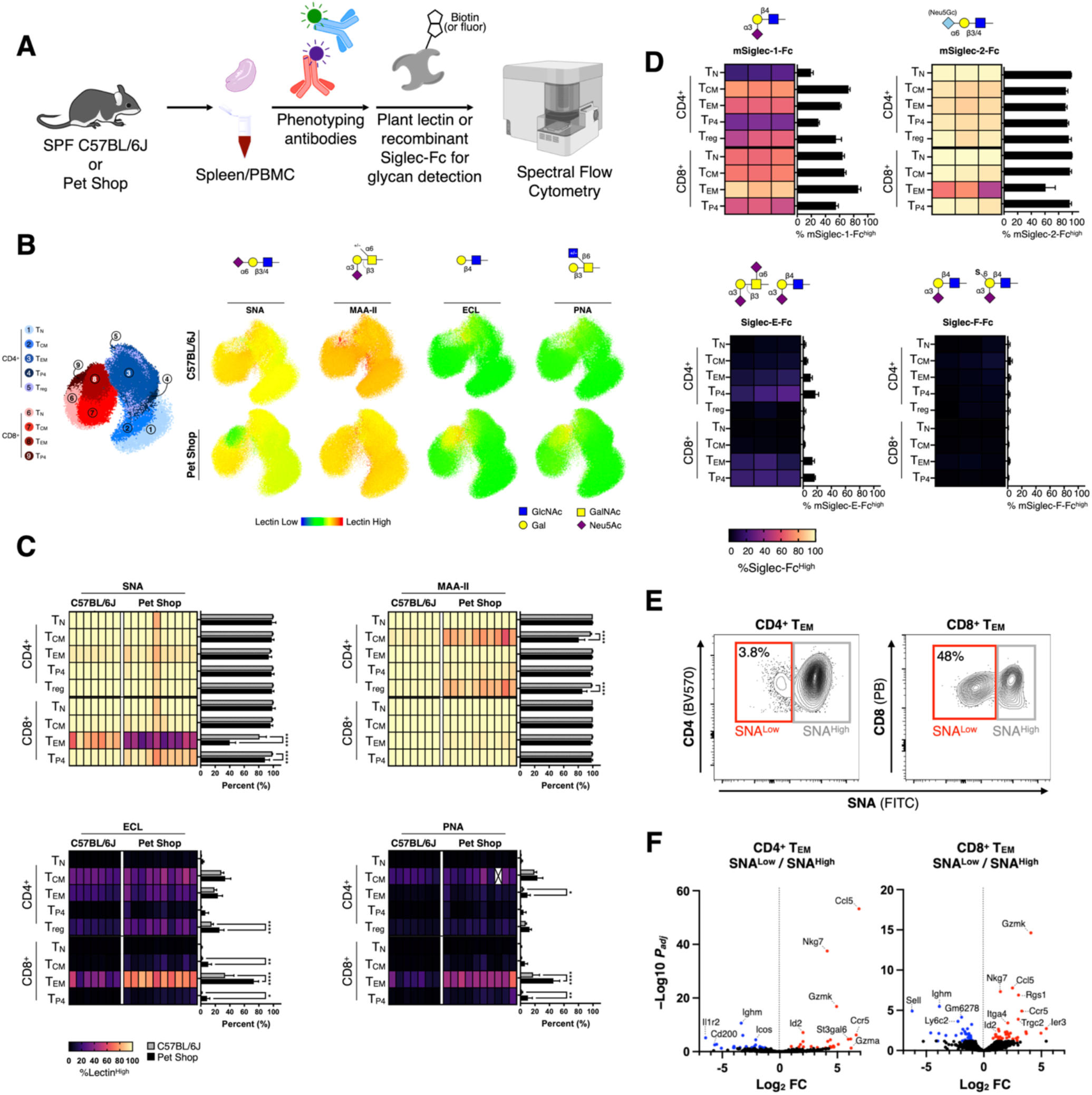
Glycocalyx composition is linked to T cell phenotype. (**A**) Glycoprofiling workflow enabled by spectral flow cytometry and glycan detecting lectin reagents. (**B**) Representative dimensionality reduced (UMAP) plots of the glycosylation profiles of splenic murine T cells from both SPF C57BL/6J and pet shop mice (6–12 weeks old). Lectin median fluorescence intensity (MFI) is overlaid as a thermal dimension. (**C**) Quantification of the percentage of each splenic T cell population that stained with each lectin from **B**. Columns represent individual biological replicates. Populations with fewer than 50 cells total were not included and are annotated by a white cell with an X. (**D**) Quantification of the percentage of each splenic T cell population from SPF C57BL/6J mice that stained with mSiglec-Fc reagents. (**E**) Representative gating scheme for purification of SNA^High^ and SNA^Low^ CD4^+^ and CD8^+^ T cells via FACS. (**F**) Differentially expressed genes from transcriptomic analysis of the purified T cell populations from **E** (n = 3). T cells were from ∼12 month old SPF C57BL/6J mice. Genes in red and blue are significantly (*P_adj_* < 0.05) up- and downregulated respectively. Mean ± SD. \**P* ≤ 0.05, \*\**P* ≤ 0.01, \*\*\*\**P* ≤ 0.0001. One-way ANOVA followed by Tukey’s multiple comparisons test. See Figure S1A for flow cytometry gating schemes. Cartoon of the spectral flow cytometer was obtained from BioRender.com.

To understand if the degree of α2-6-SG loss in T_EM_s correlated with the antigen experience of the host, we next profiled T cells from mice raised in non-SPF conditions (pet shop). These animals had significantly more T_EM_s compared to their SPF C57BL/6J counterparts, consistent with their exposure to a larger constellation of environmental antigens^39^ (**Figures S1E and S1F).** The α2-6-SG^Low^ CD8^+^ T_EM_ population from pet shop mice was significantly increased relative to other T cell populations, suggesting that antigen experience was directly correlated with glycocalyx remodelling (**Figures 1B, 1C** and **S1G**). We observed a similar phenotype in aged (12 months) SPF C57BL/6J mice, which had a T cell repertoire also strongly biased towards T_EM_s (**Figures S1E, S1F and S1H**). In contrast, CD4^+^ T_EM_s from SPF, pet shop, and aged mice showed only a modest (albeit highly significant) loss of α2-6-SGs on their surface, although ASGs were upregulated compared to CD4^+^ T_N_s (**Figures 1C**, **S1B**, and **S1C**). CD4^+^ T regulatory (T_reg_) cells also exhibited a similar increase in ASGs compared to the CD4^+^ T_N_ population, which was consistent with the dominant antigen experienced phenotype (CD44^High^ CD62L^Low^) within the T_reg_ pool and previous observations^40^ (**Figures 1C**, **S1C**, and **S1I**).

While SNA and MAA-II are useful probes for terminal sialylation motifs, they do not report on specific SG ligands for physiologically important lectins, such as the Siglec family of receptors. Most murine and human Siglecs are immunoinhibitory receptors that are expressed on the surface of various leukocytes.^20^ Engagement of Siglecs by their SGs ligands on the same cell (in *cis*) or a partner cell (in *trans*) can profoundly impact downstream immunity, making Siglecs and their ligands attractive targets for immunotherapeutic development.^21^ While Siglecs themselves are rarely expressed on T cells^41,42^, their glycan ligands are^15,43^, and can be presented for ligation in *trans* to other cells within the immune repertoire.^44^ Given our results using plant-derived lectins as probes we speculated that the relative presentation of glycan ligands for Siglecs may also be heterogeneous across different T cell populations. We tested this using four recombinant murine Siglecs fused to a human fragment crystallizable domain^43^ (mSiglec-Fc) in our spectral flow cytometry workflow (**Figure 1A**); mSiglecs-1, -2, -E, and -F. While the SG binding lectins SNA and MAA-II broadly recognize α2-6 and α2-3 sialosides respectively, Siglec-Fc reagents have more granular specificity for the core glycan structures associated with the sialic acid capping residue (glycan epitopes bound by Siglecs are illustrated in **Figure 1D**) but consequently do not report as broadly on terminal glycosylation motifs. Mutant counterparts (Arg→Ala) for each mSiglec-Fc were used as staining controls since these proteins lack the conserved arginine that is critical for SG binding.^42^ We observed striking heterogeneity in the presentation of ligands for mSiglec-1 across T cell subsets, with antigen experienced populations in both CD4^+^ and CD8^+^ compartments displaying the highest levels (**Figures 1D** and **S2A**). CD4^+^ T_N_s and T_CM_s were almost completely devoid of glycan ligands for mSiglec-1. None of the T cell populations were efficiently stained by mSiglec-E or -F. In contrast, mSiglec-2, which recognizes α2-6-SGs (with preference for those terminating in the non-human sialic acid *N*-glycolylneuraminic acid, Neu5Gc), stained cells similarly to SNA (which recognizes both *N*-acetylneuraminic acid, Neu5Ac, and Neu5Gc). Combined, plant-derived lectin and mSiglec-Fc data confirm that T cell glycosylation is heterogeneous across the major murine T cell populations that exist *in vivo*. These results suggest that murine T cells adapt their glycocalyx composition in concert with their phenotype. This further supports models^11,44^ suggesting that T cell glycocalyx remodelling promotes or disfavours interactions with cells expressing immunomodulatory lectins, such as the Siglecs (**Figure S2B**).

### α2-6-SGLow murine T cells are a highly activated effector subpopulation

While CD4^+^ and CD8^+^ T_EM_s from all animals tested adopted either an α2-6-SG^High^ or α2-6-SG^Low^glycoprofile (**Figure 1E**), these differently glycosylated populations could not be clearly delineated by the phenotyping antibodies in our panel beyond slightly increased CXCR3 expression on α2-6-SG^Low^ CD4^+^ T_EM_s (**Figure S3A**). To determine if functional and/or phenotypic differences existed between these populations, we purified α2-6-SG^High^ and ^-Low^ T_EM_s from aged mice (which had far more of the α2-6-SG^Low^ population vs. young animals) via fluorescence-activated cell sorting (FACS) and then performed bulk transcriptomic analysis (**Figures 1E** and **1F**). We observed that both CD4^+^ and CD8^+^ T_EM_s in the α2-6-SG^Low^ population were enriched in transcripts coding the effector molecules granzyme K (*Gzmk*) and chemokine ligand 5 (*Ccl5*) relative to their α2-6-SG^High^ counterparts. Increased expression of the protein products of these genes was confirmed in a separate assay (**Figure S3B**). Here, we found that even without stimulation the α2-6-SG^Low^ populations contained 2–3-fold more CCL5 and granzyme K compared to the α2-6-SG^High^ cells. Transcripts for natural killer cell granule protein 7 (*Nkg7*), which regulates cytotoxic granule exocytosis in T cells and is critical for cytolytic function^45^, were also enriched in these populations. These mRNA and protein expression data suggest that α2-6-SG^Low^ T_EM_s represent a population of highly activated T cells with elevated effector functions, including leukocyte recruitment and cytolytic capacity.

### T cell glycocalyx remodelling is driven by *de novo* glycan biosynthesis

Given that loss of α2-6-SGs was linked to an enhanced T cell effector phenotype, we next sought to dissect the mechanisms that controlled the display of these glycans on the T cell surface. We reasoned that loss of α2-6-SGs could manifest during clonal expansion of T_N_s or be a process that occurs following terminal differentiation. To test this, we stimulated murine T cells using anti-CD3/CD28 beads and measured changes in the composition of T cell glycocalyces over five daughter cell generations (**Figures 2A** and **2B**). These experiments revealed that α2-6-SGs (as detected by SNA) were lost gradually during clonal expansion of both CD4^+^ and CD8^+^ populations; a process that proceeded in concert with increased levels of corresponding ASG structures as detected by ECL and PNA (**Figures 2B** and **2C**). We also observed a modest increase in α2-3-SGs (as reported by MAA-II), but only on CD4^+^ T cells. While these data confirmed that clonal expansion was a dominant process promoting murine T cell glycan remodelling, it did not inform on the biochemical mechanisms that directly controlled these changes in SG and ASG architecture. We speculated that loss of α2-6-SGs coupled with increased ASGs could be driven by either reduced activity of the α2-6-specific sialyltransferase St6gal1^24^ during SG biosynthesis and/or catabolism of existing SGs by mammalian neuraminidase enzymes, Neu1–4^46–48^. This latter hypothesis was motivated by previous studies that showed Neu1 was presented on the surface of activated T cells and was active on an artificial reporter SG substrate.^46,48^ To test these possibilities, we stimulated T cells in the presence of chemical inhibitors of these pathways; per-*O*-acetyl-3-F(axial)-*N*-acetylneuraminic acid (3F_ax_Neu5Ac, a pan-sialyltransferase inhibitor^49,50^) or 2-deoxy-2,3-didehydro-*N*-acetylneuraminic acid (DANA, a pan-neuraminidase inhibitor that is active extracellularly^12,51^) (**Figures 2A** and **2C**). We reasoned that if St6gal1 was rate limiting in α2-6-SG biosynthesis then 3F_ax_Neu5Ac should exacerbate the decreased SNA signal that occurred during clonal expansion. Conversely, if biosynthesis remained stable but catabolism of α2-6-SGs by neuraminidases was increased then DANA should attenuate this loss. These experiments revealed that reduced sialyltransferase-mediated biosynthesis was the major driver of glycocalyx remodelling as 3F_ax_Neu5Ac significantly potentiated loss of α2-6-SGs on CD4^+^, and to a lesser extent, on CD8^+^ T cells as reported by SNA (**Figures 2C** and **2D**). Sialyltransferase inhibition also promoted a dramatic increase in asialo core 1 structures (T antigen) on CD4^+^ T cells as detected by PNA. While CD8^+^ T cells displayed increased terminal LacNAc ASGs (detected by ECL) following treatment with 3F_ax_Neu5Ac, levels of T antigen (PNA) were unchanged relative to inhibitor-naïve controls, suggesting that loss of α2-6-linked sialic acids on CD8^+^ T cells likely occurred primarily on N-linked glycans. This demonstrated that while O-linked glycan presentation was increased upon activation of CD8^+^ T cells^10,12^, the process was not due to a metabolic bottleneck in sialylation of these glycans. Inhibition of neuraminidases by DANA did not impact the composition of the glycocalyx for either T cell population, which suggested that catabolism of SGs was not the major driver of SG loss. Further, sialyltransferase inhibition through 3F_ax_Neu5Ac promoted more efficient clonal expansion of both CD4^+^ and CD8^+^ T cells under these conditions (**Figure 2E**). Thus, our data indicate that suppression of SG presentation at the biosynthetic level is the primary driver of T cell glycocalyx remodelling.

**Figure 2:**
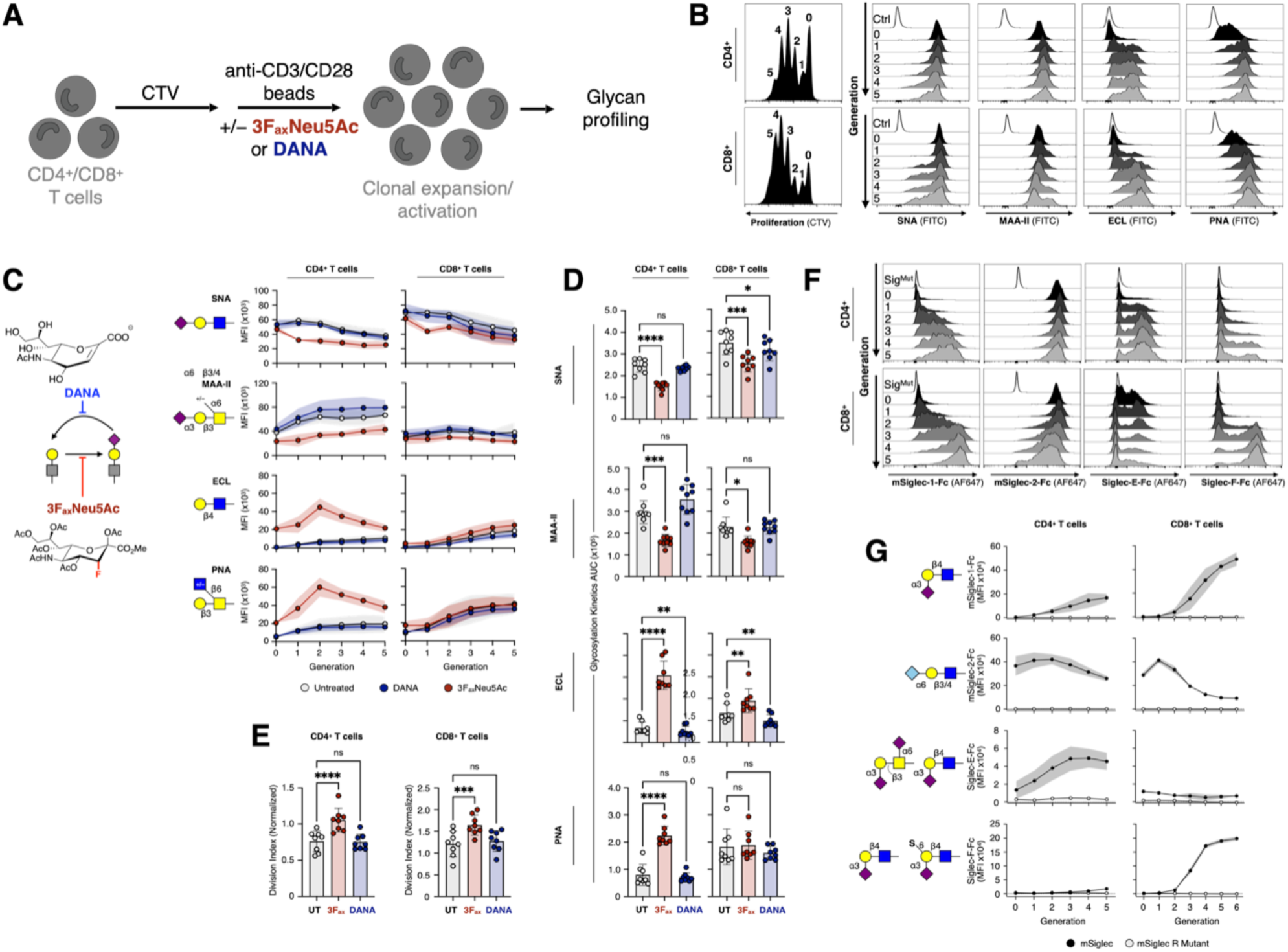
The murine T cells glycocalyx is remodelled during clonal expansion. (**A**) T cell stimulation workflow. T cells within a mixture of splenocytes from 6–12 week old SPF C57BL/6J mice were stimulated using anti-CD3/CD28 beads in the presence of 3F_ax_Neu5Ac (100 μM) or DANA (100 μM) or no inhibitor for 72 h. (**B**) Proliferation dye (cell trace violet, CTV) dilution histograms (left) and lectin signal for each T cell generation as gated on proliferation traces. (**C**) Structures and action of inhibitors of SG biosynthesis (3F_ax_Neu5Ac) and catabolism (DANA) (left) deployed in an assay measuring generational glycosylation kinetics of stimulated T cells (right). Each data point represents the MFI of a lectin for a given generation of T cell as shown in **B**. Shaded regions represent SD (N = 8). (**D**) Area under each generational glycosylation kinetic curve from **C**. 3F_ax_ = 3F_ax_Neu5Ac. (**E**) Division indices of T cells (as in **B** and **C**) as a function of inhibitor treatment. (**F**) mSiglec-Fc signal for each generation of T cell from a stimulation assay as in **A** and **B**. Shaded regions represent SD (N = 3). (**G**) Generational glycosylation kinetics as reported by mSiglec-Fc reagents from **F**. Mean ± SD. \**P* ≤ 0.05, \*\**P* ≤ 0.01, \*\*\**P* ≤ 0.001, \*\*\*\**P* ≤ 0.0001. One-way ANOVA followed by Tukey’s multiple comparisons test.

### Glycan ligands for multiple murine Siglecs are dynamically presented during clonal expansion

Glycan profiling of resting murine CD4^+^ and CD8^+^ T_N_s revealed that they presented low levels of glycan ligands for mSiglec-1, -E, and -F and higher levels for mSiglec-2 relative to other T cell populations (**Figures 1D** and **S2A**); however, these experiments did not measure potential changes in mSiglec ligand expression upon T cell stimulation. While others have demonstrated that mSiglec ligand presentation is sensitive to T cell activation^15,16,52^, these studies did not comprehensively profile both CD4^+^ and CD8^+^ compartments. Here, we found that presentation of glycan ligands for mSiglecs-1 and -F increased on each daughter cell generation for both CD4^+^ and CD8^+^ T cells after stimulation, whereas ligands for mSiglec-2 gradually decreased with a trajectory mirroring experiments using SNA (**Figures 2F** and **2G**). Ligands for Siglec-E also increased rapidly on each generation but began to recede starting in generation 3 in the CD8^+^ compartment. These results are supported by previous work showing that activated murine T cells switch sialic acid capping residues from Neu5Gc to Neu5Ac, with the latter producing superior ligands for Siglec-E.^15,16^ Our data confirms that presentation of ligands for mSiglecs in *trans* is dynamic, accompanies the early stages of murine T cell activation, and is differently regulated between the CD4^+^ and CD8^+^ compartments. Since splenic T_EM_s directly from murine spleen did not display substantial expression of ligands for Siglecs-E or -F (**Figures 1D** and **S2A**), it is likely that these SGs are downregulated on the T cell surface following clonal expansion. These data provide a dynamic dimension to known T cell-partner cell interactions that occur in *trans*; for example, with B cells through mSiglec-2^44^. In addition to known interactions, these results highlight the potential for SG-Siglec engagement between activated T cells and marginal zone macrophages through Siglec-1, neutrophils through Siglec-E, and eosinophils through Siglec-F (**Figure S2B**).^42^

### Murine T cell glycan dynamics are recapitulated *in vivo* and memory cells retain the remodelled glycocalyx

While *ex vivo* T cell activation experiments provided the specific effects of CD3/CD28 clustering on glycocalyx dynamics, they did not fully capture the array of signals received by T cells *in vivo* that may also influence glycan architecture. To account for this, we measured terminal glycosylation patterns on T cells activated in response to acute lymphocytic choriomeningitis virus (LCMV)-Armstrong (Arm) infection *in vivo*. Infection with LCMV-Arm stimulates robust antiviral T cell responses that clear infection within 8–10 days and generates memory T cells that protect against subsequent reinfection.^53^ To specifically define the evolution of glycosylation patterns by antigen-specific T cells in response to infection, we adoptively transferred LCMV-specific TCR transgenic^54^ CD4^+^ (SMARTA) and CD8^+^ (P14) T cells prior to infection (**Figure 3A**). T cell glycoprofiling eight days after infection revealed a similar glycosylation landscape to earlier experiments (**Figures 1B**–**D**), with activated T cell populations (host T_EM_s, SMARTA, and P14) exhibiting the lowest levels of α2-6-SGs and highest levels of ASG structures. Similar to experiments with anti-CD3/CD28 beads, the naïve SMARTA and P14 cells rapidly lost α2-6-SGs at early timepoints (24, 48 h) after infection (**Figure 3E**, **S4A**, and **S4B**) relative to T_N_s. This was accompanied by a major increase in ASGs as measured by ECL and PNA. While this was consistent with previous reports of increased O-linked ASGs on activated polyclonal T cells during LCMV infection (specifically T antigen detected via PNA), the effect on LacNAc presentation (detected via ECL) has not been previously described.^9,10^ We observed near-complete loss of α2-6-SGs on virus-specific CD8^+^ P14 T cells at day eight – a feature that proved durable even at extended timepoints post-clearance of LCMV (>day 14), as T cells differentiated to form a stable memory population. In contrast, virus-specific CD4^+^ SMARTA T cells eventually recovered α2-6-SGs but still retained elevated levels of ASGs in the memory phase. Host T_N_s that did not respond to viral antigen displayed minimal changes in glycocalyx composition despite the robust inflammatory response generated by LCMV infection, while the polyclonal repertoire of host T_EM_s (representing the host-derived virus-specific response) exhibited similar remodelling dynamics to their SMARTA/P14 counterparts (**Figure 3D**). These results confirm that murine T cell glycocalyx remodelling is the direct result of antigen specific activation *in vivo* and that the memory T cells retain the remodelled glycocalyx.

**Figure 3:**
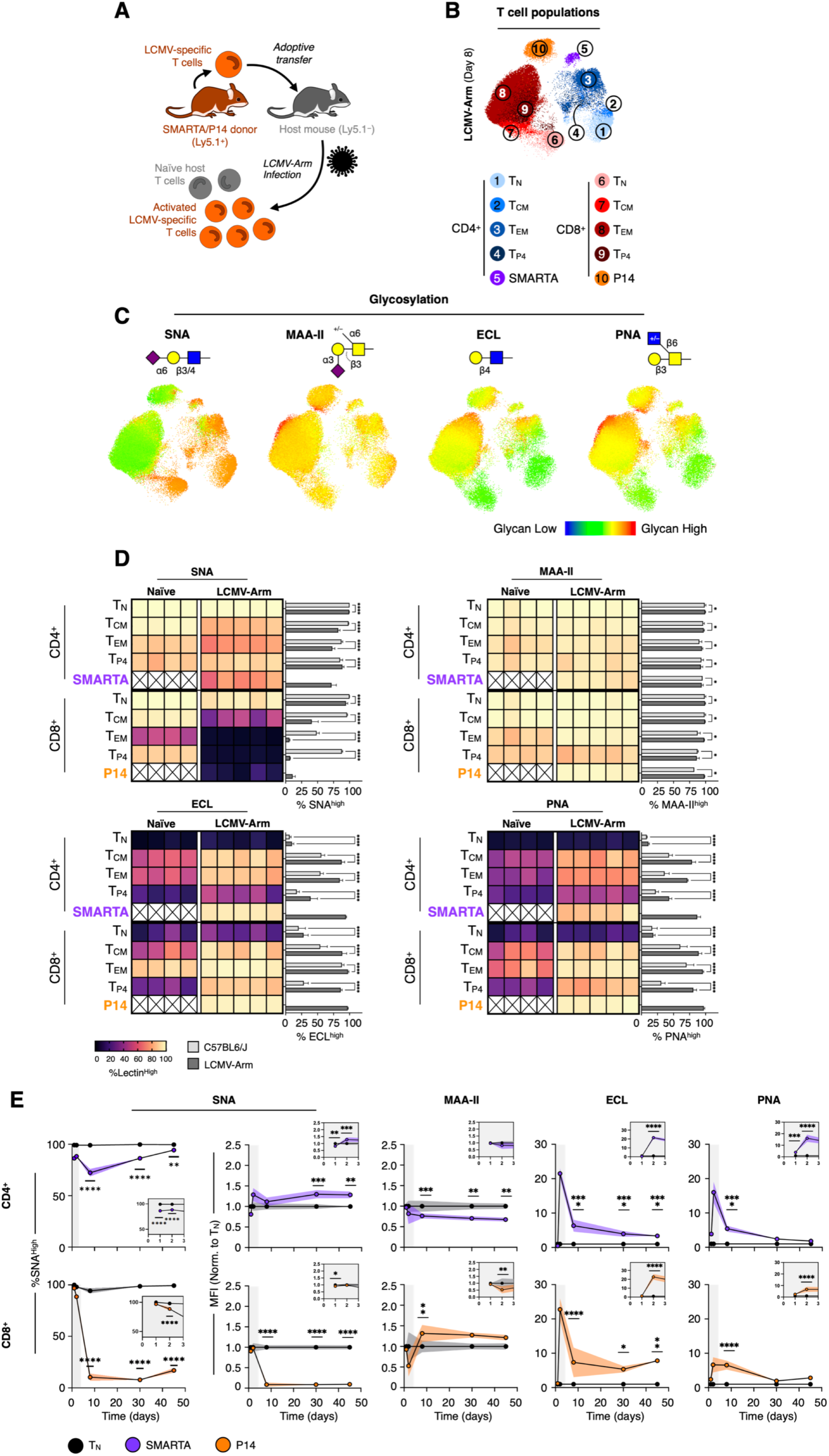
Antigen-specific activation of murine T cells *in vivo* results in glycocalyx remodelling that persists on memory cells. (**A**) Adoptive transfer and LCMV infection workflow for data in all panels. LCMV-specific CD4^+^ (SMARTA) and CD8^+^ (P14) T cells were adoptively transferred 1 day prior to acute LCMV-Arm infection. Adoptively transferred T cells are identified by Ly5.1 expression by flow cytometry. (**B**) Representative UMAP plot of the T cell repertoire of host mice 8 days following infection with LCMV-Arm. Adoptively transferred LCMV specific T cells are shown as separate clusters (SMARTA & P14, clusters 5 and 10 respectively). (**C**) Representative UMAP plots of the glycosylation profiles of T cells shown in **B**. Lectin MFI is overlaid as a thermal dimension. (**D**) Quantification of the percentage of each splenic T cell population from virus naïve or LCMV-infected mice stained with each lectin from **C**. Columns represent individual biological replicates. Naïve mice did not receive SMARTA or P14 cells and so these populations are absent as indicated by white cells with an X. (**E**) Splenic SMARTA and P14 cells were analyzed via lectin staining/flow cytometry at the indicated time points after infection. Inserts are expansions of days 1 and 2. TN = naïve T cells from host (Ly5.1^−^). Shaded regions represent SD (N = 5). Mean ± SD. \**P* ≤ 0.05, \*\**P* ≤ 0.01, \*\*\**P* ≤ 0.001, \*\*\*\**P* ≤ 0.0001. One-way ANOVA followed by Tukey’s multiple comparisons test.

### Exhausted murine T cells display a unique glycosylation profile

It has previously been shown that neuraminidase-mediated removal of SGs from co-cultures of LCMV-antigen loaded dendritic cells and PD-1^+^ exhausted CD8^+^ T cells improves T cell effector functions.^31^ However, it is unclear which terminal glycan epitopes are presented on exhausted T cells. This information is needed to understand which class(es) of SG structures are targets for neuraminidases on these cells and thus related to inhibition of T cell effector functions. To address this, we induced T cell exhaustion using two different systems – chronic viral infection and a solid tumour model. First, we profiled glycosylation of virus-specific CD4^+^ SMARTA and CD8^+^ P14 T cells 30 days after acute or chronic LCMV infection. While LCMV-Arm results in acute infection and leads to memory T cell formation, LCMV-Clone-13 infection replicates to high virus titers leading to a shift toward an immunosuppressive environment and ultimately functional T cell hyporesponsiveness (exhaustion).^55,56^ As expected, transgenic T cells exposed to LCMV-Clone-13 greatly increased expression of programmed cell death 1 (PD-1) compared to counterparts from LCMV-Arm infected animals or T_EM_s from LCMV-naïve hosts (**Figures 4A** and **4B**). P14 cells displayed a significant loss of α2-6-SGs (relative to LCMV-naïve T_EM_s) irrespective of the strain of LCMV deployed, whereas SMARTA cells retained these SGs. Interestingly, α2-3-SG presentation was similar between exhausted P14 cells and LCMV naïve T_EM_s but elevated on P14 cells exposed to LCMV-Arm. Exhausted SMARTA and P14 cells also presented significantly higher levels of ASGs compared to their counterparts from LCMV-Arm infection and LCMV naïve T_EM_s. Given that translational interest in T cell exhaustion is often focused on cancer immunotherapy, we next investigated if our findings from LCMV infection were recapitulated in a solid tumour model (pancreatic ductal adenocarcinoma, PDAC^57,58^). The majority of infiltrating CD4^+^ and CD8^+^ T_EM_s from PDAC tumours established on C57BL/6J mice expressed high levels of PD-1 (**Figures 4C**), unlike splenic T_EM_s from untreated animals (**Figures 4A**, **4B**, and **S4C**). The glycosylation profiles of PDAC infiltrating T_EM_s were largely in alignment with exhausted SMARTA and P14 counterparts from the chronic LCMV model: CD8^+^ T_EM_s had lost nearly all α2-6-SGs whereas CD4^+^ T_EM_s retained these glycans and α2-3-SGs were similarly presented on both T cell subsets. Combined, results from both viral infection and tumour models confirmed that exhausted CD8^+^ murine T cells were deficient in α2-6-SGs, displayed increased levels of ASGs compared to memory CD8^+^ T cells (30 days after LCMV-Arm infection), and presented α2-3-SGs similarly to non-exhausted T_EM_ counterparts (i.e. splenic PD-1^−^T_EM_s). These data confirm that exhausted murine T cells have a combination of terminal glycan epitopes that are distinct from non-exhausted effectors and memory T cells.

**Figure 4:**
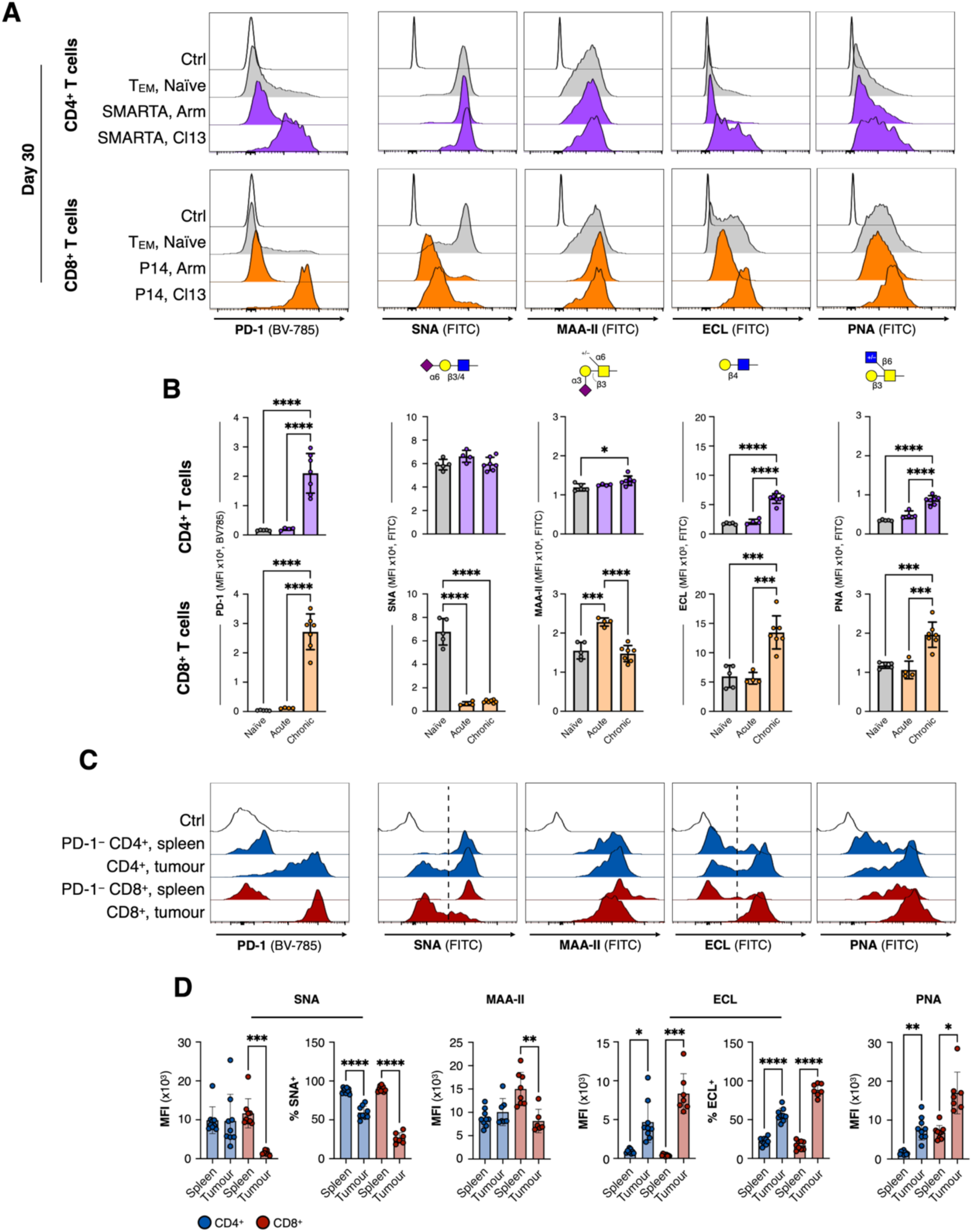
Exhausted murine T cells are uniquely glycosylated. (**A**) Representative glycosylation profiles of polyclonal T_EM_s from LCMV-naïve mice vs. LCMV-specific SMARTA and P14 T cells from mice infected with either acute LCMV-Arm or chronic LCMV-Clone 13. Flow cytometry analysis performed at day 30 after LCMV infection. Open black histograms represent lectin staining controls. (**B**) Quantification of data from **A**. (**C**) Representative glycosylation profiles of splenic PD-1^−^ and tumour infiltrating T cells from mice bearing PDAC tumours. (**D**) Quantification of data from **C**. SNA data is reported in two ways due to the bimodal distributions of the signals: left: MFI describing absolute signal intensity and right: the % of cells that stain positively with SNA. Error bars represent SD (N = 5). Mean ± SD. \**P* ≤ 0.05, \*\**P* ≤ 0.01, \*\*\**P* ≤ 0.001, \*\*\*\**P* ≤ 0.0001. One-way ANOVA followed by Tukey’s multiple comparisons test.

### T cell glycan remodelling is driven by altered expression of multiple glycosyltransferases

Experiments using 3F_ax_Neu5Ac showed that altered glycan biosynthesis was the primary driver of glycocalyx remodelling on murine T cells following *ex vivo* activation (**Figures 2C** and **2D**); however, they did not directly identify the specific biosynthetic pathways that were altered during T cell activation. To address this, we performed differential expression analysis of glycan-associated genes from scRNA-seq data from two published studies^59,60^ analyzing virus-specific CD4^+^ and CD8^+^ T cells isolated on day ten following acute LCMV-Arm infection. These virus-specific cells were compared to CD4^+^ or CD8^+^ T_N_s from the same hosts (**Figures 5A** and **5B**). Transcription of genes coding for a wide range of proteins involved in glycan biosynthesis were differentially regulated following exposure to antigen. In particular, transcripts coding for enzymes involved in the early stages of N-linked glycan biosynthesis were upregulated, including components of the oligosaccharyltransferase complex (**Figure 5C**) and several α1,3-mannosyl-glycoprotein 2-β-*N*-acetylglyucosaminyltransferases (*Mgat1*, *2*, and *4*), which control N-linked glycan branching and are known to negatively regulate T cell activation (**Figure 5D**).^14,61–63^ In our lectin assays, this manifested as increased binding of the branched N-glycan specific lectins PHA-L and PHA-E (*Phaseolus vulgaris* erythroagglutinin and leucoagglutinin, respectively) on T_EM_s compared to T_N_s (**Figure S5A and S5B**). Binding of the mannose-specific lectins Concanavalin A (ConA; from *Canavalia ensiformis*) and GNL (from *Galanthus nivalis*) suggested that only minor changes in mannose-terminating glycans manifested on CD4^+^ T_EM_s, although T_CM_ and T_P4_ populations exhibited more significant differences vs T_N_s in both CD4^+^ and CD8^+^ compartments (**Figure S5C and S5D**). O-linked glycan biosynthesis was also impacted, predominantly in construction of core 1 and core 2 structures, consistent with increased PNA staining on T_EM_s vs T_N_s (**Figures 1C** and **5E**). Importantly, major changes in sialyltransferase expression were detected, with transcripts coding for S6tgal1 – the dominant enzyme that installs sialic acid in the α2-6 configuration on N-linked glycans – significantly downregulated (**Figure 5F**). *St6gal1* was more dramatically downregulated in CD8^+^ T cells vs CD4^+^ counterparts (fold change ∼500 and 8 respectively), consistent with our observations of more rapid and durable loss of α2-6-SGs on activated CD8^+^ T cells as measured by SNA (**Figures 2C** and **3E**). In contrast, transcripts coding for several α2-3-specific sialyltransferases were upregulated, including *St3gal1*, *4*, and *6*, which act on both N- and O-linked glycans. These transcriptomic observations were in alignment with our lectin staining data which showed increased α2-3-sialylation following activation of both CD4^+^ and CD8^+^ T cells as reported by MAA-II (**Figures 2C**, **2G**, and **3E**). Changes in glycosaminoglycan and ganglioside biosynthesis (**Figures 5G and 5H**) and N-linked glycan catabolism (**Figure 5I**) were also highlighted in the analysis. Further, expression of genes related to nucleotide sugar assembly was impacted (**Figure 5I**), with global upregulation of pathways that control the biosynthesis of: UDP-galactose, -glucose, -*N*-acetylgalactosamine, -glucouronic acid; GDP-mannose; and CMP-sialic acid. Combined, lectin staining, and transcriptomic analysis provide a comprehensive overview of the changes in glycan biosynthesis that occur upon stimulation of murine T cells. Importantly, the changes in glycosylation we observed via lectin staining mapped well onto the known functions of glycosyltransferases that were found to be differentially regulated. The results suggest that murine T cells control glycocalyx remodelling upon activation through careful tuning of glycosyltransferase expression at different stages of glycan biosynthesis.

**Figure 5:**
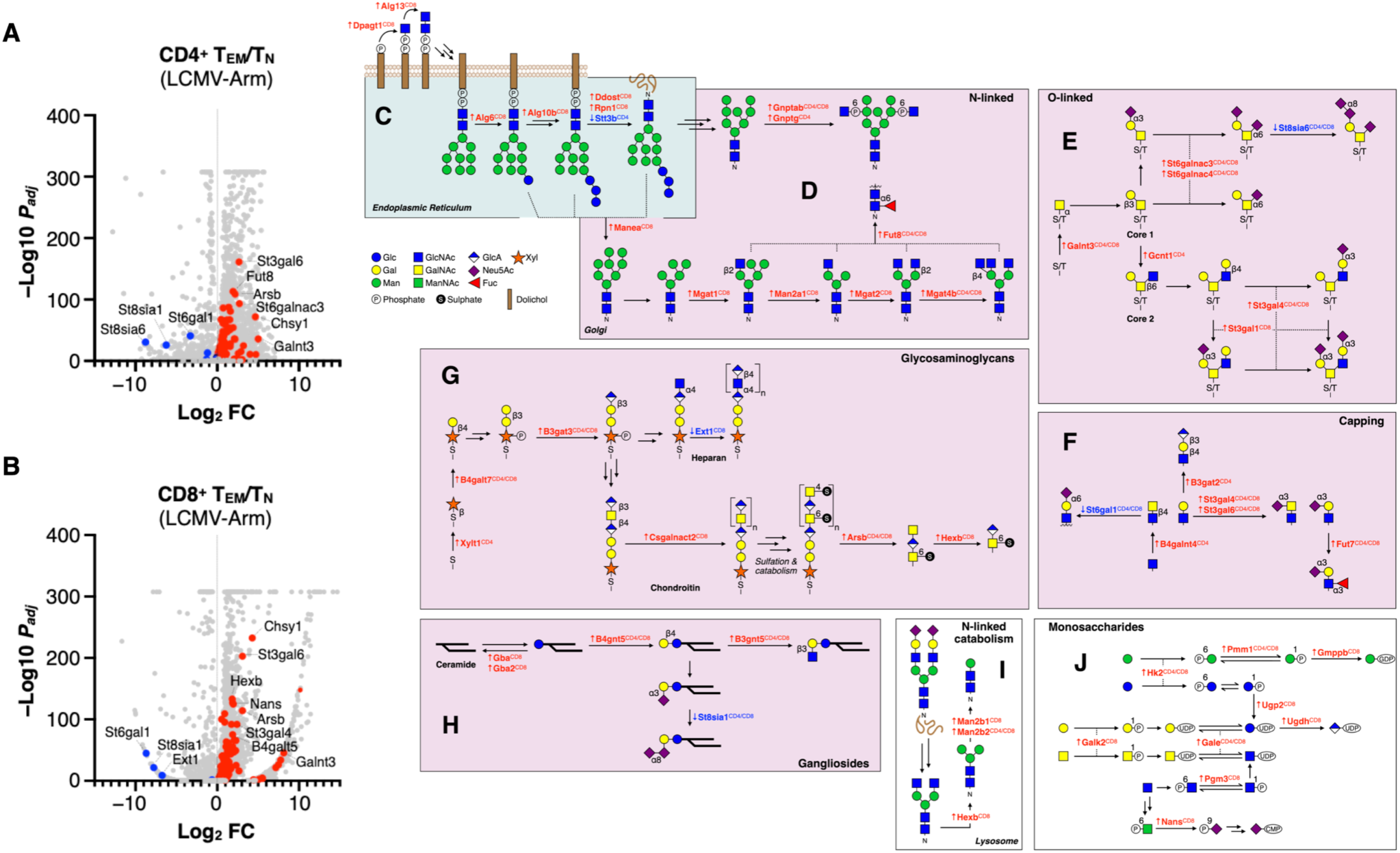
Glycan biosynthesis pathways that drive murine T cell glycocalyx remodelling. (**A**) Differentially expressed genes between LCMV-specific murine CD4^+^ T_EM_s and T_N_s. Genes in red and blue are significantly (*P_adj_* < 0.05) upregulated and downregulated respectively. Original dataset from references^59,60^. (**B**) Same analysis as in **A** but for CD8^+^ T_EM_ and T_N_. (**C**–**J**) Biosynthetic pathway schematics. Protein products of select genes that are up- (red) or down- (blue) regulated from **A** and **B** are listed above enzymatic reaction arrows. The T cell subset(s) (CD4^+^ or CD8^+^) in which the gene is differently expressed is indicated in superscript. Schematic adapted from the GlycoMaple database^71^. (**C**) N-linked glycan biosynthesis in the ER. (**D**) N-linked glycan mannose trimming and branching. (**E**) O-linked glycan biosynthesis. (**F**) Capping of complex-type N-linked glycans. (**G**) Glycosaminoglycan biosynthesis. (**H**) Ganglioside biosynthesis. (**I**) Lysosomal catabolism of N-linked glycans. (**J**) Monosaccharide metabolism and nucleotide sugar biosynthesis.

### T cell glycocalyx remodelling is divergent between mouse and human

Having characterized glycan presentation and dynamics across the murine T cell repertoire, we next applied a similar profiling strategy to human T cells. Our first experiments focused on glycan profiling of peripheral blood T cells from adults (**Figures 6A**, and **6B**, and **S6A**). We were surprised to discover that, unlike mouse, all human T cell populations from all donors were near-homogeneously glycosylated with respect to both SG and ASG structures, including T_N_ (CCR7^High^ CD45RA^High^), T_EM_ (CCR7^Low^ CD45RA^Low^), T_CM_ (CCR7^High^ CD45RA^Low^, and T_EMRA_ (CCR7^Low^ CD45RA^High^) populations. Further, all human T cell subsets presented substantial terminal LacNAc ASGs as reported by high ECL but low PNA staining, unlike murine counterparts which were broadly deficient in the ASGs recognized by both lectins at steady state (**Figures 1B** and **1C**). Our results in mouse were collected using T cells from 6–12 week old animals, and so we speculated that age may be an uncontrolled variable that could explain the differences in T cell glycosylation between the species. We addressed this in two ways: profiling T cells from (1) aged mice (1 year old) and (2) human umbilical cord blood. Aged SPF C57BL/6J mice displayed a T cell repertoire that closely resembled their outbred pet shop counterparts, with substantially more T_EM_s adopting the α2-6-SG^Low^ phenotype vs. young SPF animals (**Figures S1E**). This suggested that antigen experience, and not age, was the primary driver of T cell glycocalyx remodelling in mice since both pet shop and aged SPF mice had more opportunities for antigen exposure vs. younger SPF counterparts as measured by the relative abundance of splenic T_EM_s (**Figure S1E and S1F**). Next, we performed glycoprofiling of T cells from human cord blood, which contained mostly fetal T_N_s. We reasoned that these cells would represent a superior comparison vs. adult human PBMCs to T cells from young C57BL/6J mice. We found that a small percentage – ∼1–5% – of both CD4^+^ and CD8^+^ T_N_s from human cord blood adopted the α2-6-SG^Low^ phenotype (**Figures 6B**–**E**), whereas α2-6-SG^Low^ T cells were almost completely absent from the adult repertoire. While this result revealed that α2-6-SG^Low^ T cells exist in humans, no corresponding population was detected in the murine T_N_ compartment (**Figures 1C** and **S1B**). The differences were not tissue-dependent since T cells from murine peripheral blood (**Figure S1D**) and spleen (**Figure 1C**) were similarly glycosylated and neither resembled T cells from human peripheral or cord blood. We next postulated that human T cells with diminished α2-6-SG presentation may not manifest in the healthy adult donor repertoire without recent stimulation. To test this, we stimulated human T cells in a generational glycosylation kinetic profiling assay using an approach similar to experiments with murine counterparts (**Figures 2A**, **7A–C**). These experiments revealed a striking divergence in glycocalyx dynamics as a function of clonal expansion between the species as α2-6-SGs greatly increased on the first daughter cell generation and remained elevated in subsequent generations (**Figure 7B**). This result contrasted with experiments using murine T cells where the same α2-6-SGs gradually diminished as clonal expansion proceeded (**Figures 2B** and **2E**). Importantly, CD4^+^ T cells that had not divided were ∼60% antigen experienced (CD45RA^Low^) and were not capable of clonally expanding under the conditions tested (**Figures 7D** and **7E**). These CD45RA^Low^ T cells retained intermediate α2-6-SG presentation even in the presence of anti-CD3/CD28 beads. This result suggests that at an early stage following TCR activation, human CD4^+^ T cells increase presentation of α2-6-SGs but later lose these glycans as they enter a more terminally differentiated state. Similar experiments using human Siglec-Fc reagents (hSiglec-Fc) revealed that both CD4^+^ and CD8^+^ populations displayed chronically high presentation of ligands for hSiglecs-1, -2, and -7, which recognize α2-3-, α2-6-, and α2-8-SGs/disialyl T antigen, respectively (**Figures S6B** and **S6C**). The presentation of Siglec ligands was only modestly affected by T cell stimulation and only for some of the PBMC donors. However, clonal expansion did significantly reduce the number of CD4^+^ T cells with the highest Siglec-7 ligand density, which was consistent with observations made by others under different activation conditions.^64^ Thus, human Siglec ligand dynamics are divergent from those in mice since murine T_N_s presented relatively low levels of multiple mSiglec ligands until activated (**Figure 2G**). Our observations suggest that human T cells retain the capacity to interact with other immune cells through Siglec:glycan interactions in *trans* irrespective of stimulation. These results point towards fundamental differences in the dynamics of glycan-level regulation of T cell immune responses between the species, and critically, confirm that divergence in T cell glycosylation between mouse and human is a *bona fide* fundamental physiological difference and not directly related to age, antigen experience, or tissue of origin.

**Figure 6:**
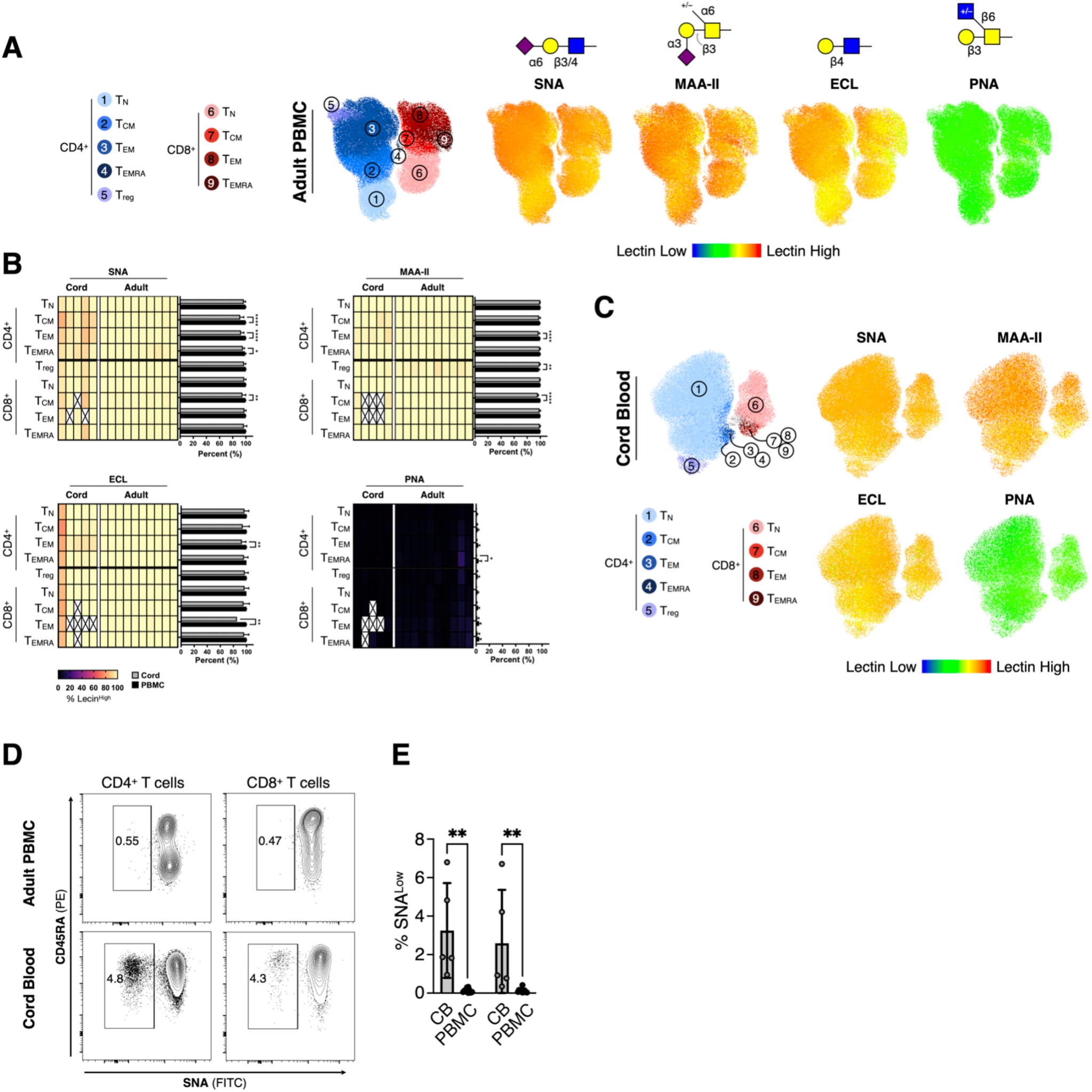
Human T cell glycocalyces are not linked to specific phenotypes at rest. (**A**) Representative dimensionality reduced (UMAP) plots of the glycosylation profiles of healthy adult (20–30 years) human peripheral blood T cells. Lectin signal is overlaid as a thermal dimension. T_EMRA_ = T effector memory CD45RA^High^ cells. (**B**) Quantification of the percentage of each T cell population that stained with each lectin from **B** and **C**. Columns represent individual biological replicates. Populations with fewer than 50 cells total were not included and are annotated by a white cell with an X. (**C**) Analysis of the T cell repertoire from human cord blood analyzed as in **A**. (**D**) Representative flow cytometry density maps of T cells from human cord blood specifically highlighting the small SNA^Low^ population. (**E**) Quantification of data from **D** (N = 5). Mean ± SD. \*\**P* ≤ 0.01. One-way ANOVA followed by Tukey’s multiple comparisons test.

**Figure 7:**
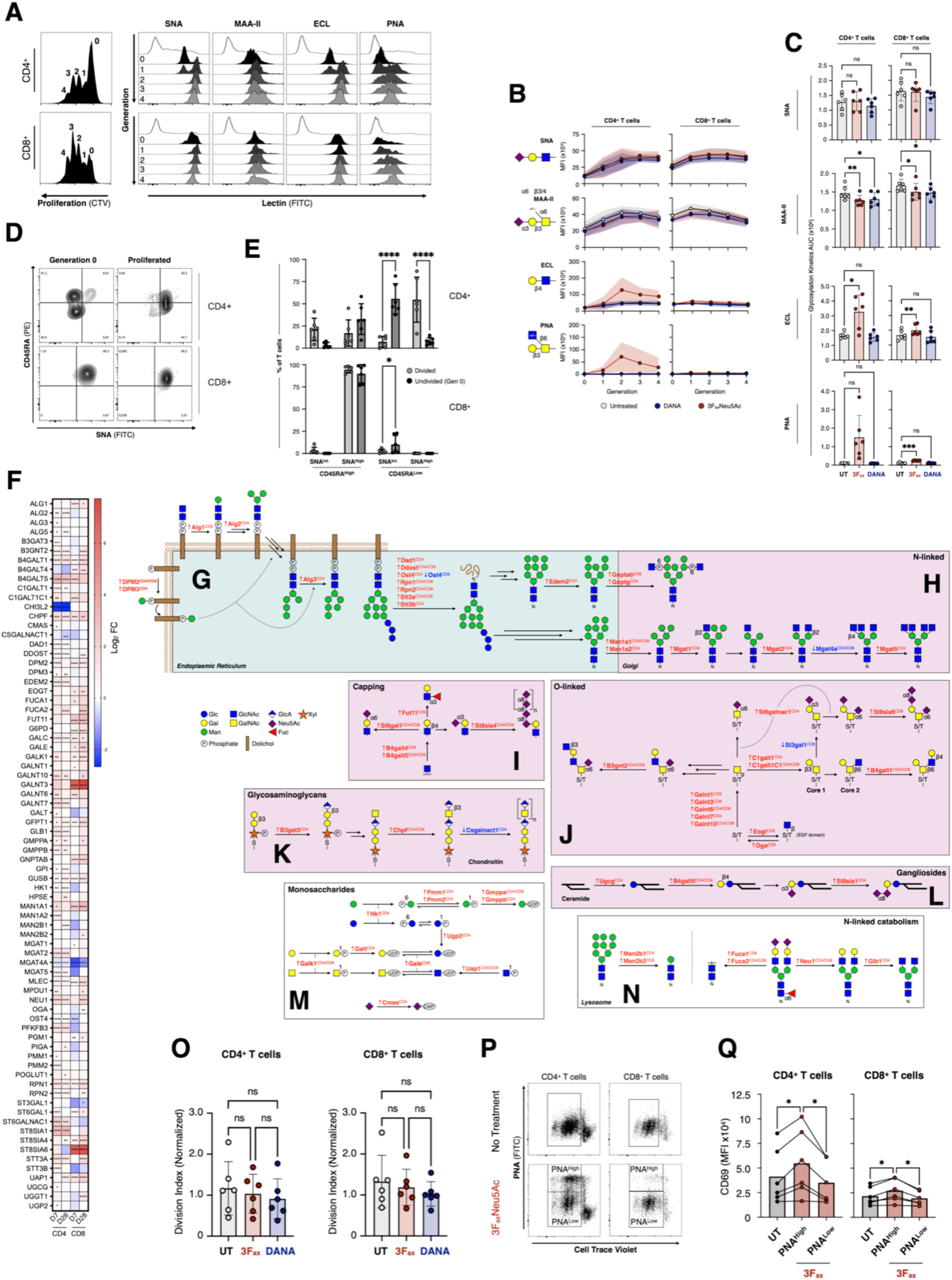
Human T cell glycocalyx remodelling occurs during clonal expansion but in manner distinct from murine counterparts. (**A**) Human T cells within a mixture of PBMCs from 20–30 year old healthy adults were stimulated using anti-CD3/CD28 beads for 72 h at 37 °C. Proliferation dye (cell trace violet, CTV) dilution histograms (left) and lectin signal for each T cell generation as gated on proliferation traces. (**B**) Generational glycosylation kinetics of stimulated human T cells (as in **A**) cultured with 3F_ax_Neu5Ac (100 μM) or DANA (100 μM) or no inhibitor for 72 h. Each data point represents the median fluorescence intensity (MFI) of a lectin for a given generation of T cell (as in **A**). Shaded regions represent SD (N = 6). 3F_ax_ = 3F_ax_Neu5Ac. (**C**) Area under each generational glycosylation kinetic curve from **B**. (**D**) Flow cytometry density maps showing the enrichment of CD45RA^Low^ cells in the undivided (generation 0) population (see **A**). (**E**) Quantification of data from **D**. SNA^Int.^ = intermediate SNA signal. (**F**) Significantly (*P_adj_* < 0.05) differentially expressed glycan processing genes between CD4^+^ or CD8^+^ T_EM_ and T_N_ from heathy adult humans infected with SARS-CoV-2 (see reference^65^). Only genes with fold changes of ≥ 2 or ≤ −2 are included. D7 and D28 = day 7 and day 28 post-infection respectively. (**G**–**N**) Biosynthetic pathway schematics. Protein products of select genes that are up- (red) or down- (blue) regulated from **A** and **B** are listed above enzymatic reaction arrows. The T cell subset(s) (CD4^+^ or CD8^+^) in which the gene is differently expressed is indicated in superscript. Schematic adapted from the GlycoMaple database^71^. (**G**) N-linked glycan biosynthesis in the ER. (**H**) N-linked glycan mannose trimming and branching. (**I**) Capping of complex-type N-linked glycans. (**J**) O-linked glycan biosynthesis. (**K**) Glycosaminoglycan biosynthesis. (**L**) Ganglioside biosynthesis. (**M**) Monosaccharide metabolism and nucleotide sugar biosynthesis. (**N**) Lysosomal catabolism of N-linked glycans. (**O**) Division indices of T cells (as in **A** and **B**) as a function of inhibitor treatment. (**P**) Gating scheme for identifying T cells maximally impacted by 3F_ax_Neu5Ac (PNA^High^) as in **B**. (**Q**) Quantification of CD69 expression levels as measured by flow cytometry on untreated (UT) T cells or counterparts exposed to 3F_ax_Neu5Ac (3F_ax_), which are subdivided into PNA^Low^ and PNA^High^ populations as in **P**. The outcome for each patient sample is shown via connecting lines. Mean ± SD. \**P* ≤ 0.05, \*\**P* ≤ 0.01, \*\*\**P* ≤ 0.001, \*\*\*\**P* ≤ 0.0001. One-way ANOVA followed by Tukey’s multiple comparisons test.

### Human T cell glycan biosynthesis is regulated by antigen experience

We next sought to understand if species differences in T cell glycosylation could be explained by divergent regulation of glycan biosynthesis. To investigate this, we turned to recently published single-cell transcriptomic datasets comparing gene expression profiles of human peripheral blood T cells from healthy individuals before and after deliberate infection with an early strain (pre-Alpha) of SARS-CoV-2, which induces robust T cell activation in humans.^65^ Our rationale was that these data would be most comparable to T cells from LCMV-infected mice (**Figure 5**) because both datasets involved viral infection and contained naïve, ∼1-week, and ∼1-month post-infection treatment groups. As with our analysis of murine transcriptomic data, we focused exclusively on genes known to be related to glycan processing (**Figure 7F**). Here we found that activated human T cells also upregulated genes involved in the early stages of N-linked glycan assembly, including many oligosaccharyltransferase (OST) complex-related proteins and the mannosidase enzymes MAN1A1 and 2 (**Figures 7G** and **7H**). MGAT2 and 5 were upregulated in CD4^+^ T cells but MGAT4 was downregulated in both CD4^+^ and CD8^+^ populations, suggesting more granular regulation of N-linked glycan branching in human T cells. Upregulation of these MGATs was further supported by our observation that PHA-E and PHA-L staining increased on T_EM_s relative to T_N_s, which suggested potentiated N-linked glycan branching on T_EM_s (**Figures S5E and S5F**). Staining with ConA revealed that human T_EM_s presented significantly more terminal high-mannose structures (**Figure S5G**), which suggested that while N-linked glycan biosynthesis was likely enhanced in activated human T cells, downstream complex-type processing steps were unable to accommodate the increased metabolic load. We did not detect major differences in GNL staining between any human T cell populations, which contrasted with results from mouse T cells (**Figure S5D** and **S5H**). Even so, transcripts coding for two galactosyltransferases (B4GALT4 & 5) that control LacNAc biosynthesis on complex-type glycans^66^ were found to be upregulated in both CD4^+^ and CD8^+^ T_EM_s and this change clearly manifested as increased binding of ECL to stimulated T cells (**Figures 7A** and **7I**).

O-linked glycan biosynthesis was also significantly altered in activated human T cells, with expression of many functionally overlapping *N*-acetylgalactosamine transferases (GALNT family, install GalNAc as the first step in core 1 and 2 biosynthesis^67^) increased (**Figure 7J**). To test if differential regulation of GALNTs manifested as altered glycosylation, we measured α*O*-GalNAc (Tn antigen) on activated T cells using the Tn-binding lectin *Helix aspersa* agglutinin (HAA^37^) (**Figure S6D**). We found that HAA signal was increased on activated T cells, which was consistent with increased production of Tn antigen by GALNTs and previous observations.^68^ Increased elaboration of Tn antigen to core 1 (T antigen) by upregulation of the core 1 synthase glycoprotein-*N*-acetylgalactosamine 3-β-galactosyltransferase 1 (C1GALT1^69^) was also confirmed via PNA staining (**Figure 7A**). In addition, we found that the early stages of chondroitin biosynthesis and ganglioside assembly were upregulated (**Figures 7K** and **7L**), in addition to pathways controlling the biosynthesis of most major classes of nucleotide sugars and N-linked glycan catabolism (**Figures 7M** and 7**N**).

Genes coding for multiple sialyltransferases were upregulated across both CD4^+^ and CD8^+^ subsets (**Figures 7I** and **7J**) including *ST6GAL1*, which was in contrast, downregulated in murine T cells (**Figure 5F**). This observation explained the striking divergence in α2-6-sialylation trajectories between the species (**Figures 2B**, **2C**, **2G**, **7A**, and **7B**). Importantly, expression of *ST6GAL1* returned to T_N_ levels ∼one month post-infection, which was consistent with our observation that α2-6-SGs on antigen experienced human T cells decreased to match levels on other T cell populations over the long term (**Figures 6A**, **6B**, and **7F**). The only gene coding for a sialyltransferase found to be downregulated was *ST3GAL1*, which was consistent with the known changes in sialylation of core-1 O-linked glycans on activated murine T cells.^22^ Even with diminished *ST3GAL1*, we did not detect major changes in global α2-3-SG presentation upon stimulation of human T cells (**Figure 7A**).

To more deeply probe the impact of these gene expression changes on glycan presentation, we repeated experiments using 3F_ax_Neu5Ac (see **Figure 2A**) using human T cells (**Figures 7B** and **7C**). We found that 3F_ax_Neu5Ac did not impact biosynthesis of α2-6-SGs as reported by SNA (unlike experiments with murine T cells, **Figures 2C** and **2D**); however, a small decrease in MAA-II signal was observed (**Figure 7C**), which suggested that some α2-3-sialylation events were compromised. An explanation for these results is that upregulation of ST6GAL1 in human T cells made α2-6-sialic acid capping events less susceptible to chemical inhibition whereas downregulation of ST3GAL1 had the opposite effect. 3F_ax_Neu5Ac also potentiated presentation of ASGs on human T cells as reported by ECL and PNA, which was likely the result of preventing ST3GAL1-mediated α2-3-sialylation of O-linked glycans. Unlike experiments with murine T cells (**Figure 2E**), sialytransferase inhibition did not augment clonal expansion (**Figure 7O**); however, it did significantly increase expression of the early T cell activation marker CD69 on both T cell populations (**Figures 7P** and **7Q**). The effect was only observed for the T cells that were maximally impacted by sialyltransferase inhibition (PNA^High^ cells) (**Figure 7P**) and is consistent with phenotypic changes that result following neuraminidase treatment.^34^ As with studies using murine T cells, inhibition of neuraminidases using DANA did not have a significant effect on glycosylation at any generation (**Figures 7B** and **7C**). Combined, these data add mechanistic context for our lectin staining results and provide further motivation to target glycan-metabolism for controlling the phenotype and function of human T cells.

## DISCUSSION

We comprehensively characterized the glycosylation profiles of major mouse and human T cell populations using spectral flow cytometry, lectins with defined glycan ligand specificities, and transcriptomic approaches. Our results show that murine T cell glycosylation is tied to phenotype at steady state, with small populations of T_EM_s from young SPF C57BL/6J mice displaying low amounts of α2-6-SGs and increased corresponding ASG structures. We used multiple mouse models (pet shop, aged, and viral infection) to confirm that the relative abundance of this alternatively glycosylated population increased with the antigen experience of the mouse. We discovered that both CD4^+^ and CD8^+^ α2-6-SG^Low^ murine T cells had increased production of granzyme K and CCL5 over other populations, suggesting these cells were some of the most activated effectors in the T cell repertoire. Stimulation of murine T_N_ both *in vitro* and *in vivo* resulted in rapid loss of α2-6-SGs, increased ASGs, and dramatic remodelling of glycan ligands for multiple immunoregulatory mSiglecs. We used chemical probes and transcriptomic datasets to confirm that these changes were primarily the result of altered glycan biosynthesis rather than catabolism of SGs.

Our analysis of the human T cell repertoire using similar experimental approaches revealed major differences compared to mouse T cells. We found that human T cells from adult peripheral blood were near-homogeneously glycosylated across functionally distinct populations, whereas a small percentage of T_N_s from fetal tissue were bimodal for α2-6-sialylation. Upon stimulation, clonally expanding adult human T_N_s rapidly accumulated α2-6-SGs, which was in direct contrast to observations from murine experiments. Stimulated T cells eventually reduced presentation of these glycans and asialo structures as they transitioned to T_EM_ and T_CM_ phenotypes. As with mouse, human T cell glycosylation dynamics could be traced back to specific biosynthetic pathways by cross-referencing lectin staining and chemical probe results with transcriptomic datasets.

These results provide context for previous observations linking T cell glycosylation to regulation of activation and effector functions. Both mouse^28,31^ and human^34,35,48^ T cells activate more efficiently in the presence of neuraminidases, suggesting that there is a conserved SG motif that suppresses their capacity to clonally expand and deploy effector cytokines. While changes in sialylation on human CD45RA/CD45RO have been specifically measured in previous work^12,70^, it remains unclear if broader changes in glycosylation occur on other proteins that are important to T cell immunophysiology. Since our results show that activated murine T cells are deficient in α2-6-SGs but human counterparts upregulate these structures, it is unlikely that α2-6-SGs are the functionally important targets for T cell stimulating neuraminidases. Instead, our results implicate α2-3-SGs as T cell-intrinsic immunosuppressive glycans given that they are stably presented on all major T cell subsets from both species irrespective of stimulation conditions. This is further supported by the observation that an α2-3-specific neuraminidase (sialidase from *Streptococcus pneumoniae*, SS) can enhance clonal expansion of murine T cells in *ex vivo* stimulation assays^31^, much like pan-linkage selective counterparts which have maximal activity on α2-3-linked structures.^18^ We verified that SS is capable of efficiently removing α2-3-linked sialic acids from SGs on all T cell subsets from both mouse and human (**Figure S7**).

These results are of particular significance to recent efforts focused on harnessing neuraminidase-based technologies to stimulate beneficial anti-tumour immune responses^34,35^, which have been developed partially in murine models. They are also complementary to findings related to N-linked glycan branching on T cells – a process that can be manipulated to negatively regulate TCR signalling for therapeutic benefit in settings of autoimmunity.^8,14,61–63^ With this cross-species atlas of T cell glycophysiology in hand, next-generation glycan-targeted therapeutics can be designed for specific activity on functionally important glycans on human T cells – for example, α2-3-SGs – to ensure maximal translational impact.

## METHODS AND SUPPLEMENTAL INFORMATION

Methods are included at the end of this manuscript and supplemental information can be found in a companion document.

## ACKNOWLEDGEMENTS

We thank Sylvia Okafor, Candaice Newell, Fariba Baghai Wadji, Sharon Miksys, Wai Haung (Ho) Yu, and Samuel P. Nyandwi for assistance with mouse tissue collection. Flow cytometry assistance from Parva Thakker and Vincent Cheng was greatly appreciated. We gratefully acknowledge Dr. Paola Cappello for the generous donation of CR705 cells for the PDAC model. We thank Tania Watts, Arthur Mortha, Goetz Ehrhardt, Jennifer Gommerman, and Shannon Dunn for productive discussions and Andrew Thompson and Mark Nitz for critical comments on the manuscript. H. Choksi and V. Affe are grateful for scholarship support from The Canadian Institutes of Health Research (CIHR). F. N. Izzati is grateful for scholarship support from the Indonesian Endowment Fund for Education (LPDP) and support from the National Research and Innovation Agency of the Republic of Indonesia. This research was supported by grants from the following agencies to L. J. Edgar: CIHR (PTT-190383), New Frontiers in Research Fund (NFRFE-2022-00237), The Arthritis Society of Canada (IIG-22-0000000126), The Canadian Glycomics Network (CR-22), and Natural Sciences and Engineering Research Council of Canada (RGPIN-2022-04437, for chemical synthesis support). D. G. Brooks, J. Matthews, and H. Cui acknowledge support from CIHR (PJT-505111 and FDN-353612 to DGB; PTJ-162160 to JM; PJT-190073 to HC). DGB also acknowledges support from the National Institutes of Health (AI085043).

## AUTHOR CONTRIBUTIONS

L. J. Edgar, F. N. Izzati and H. Choksi designed the study. F. N. Izzati and H. Choksi performed functional immunoassays, lectin staining, and flow cytometry analysis of mouse and human tissues. P. Giuliana performed lectin staining and flow cytometry analysis of PDAC tissues and provided flow cytometry support for analysis of human samples. D. Abd-Rabbo (advised by D. G. Brooks) performed bioinformatics analysis of single-cell transcriptomic datasets. H. Elsaesser (advised by D. G. Brooks) conducted TCR transgenic T cell adoptive transfers and LCMV infections. A. Blundell chemically synthesized 3F_ax_Neu5Ac and DANA. V. Affe performed FACS purification and bulk transcriptional analysis of murine T cells. V. Kannen (advised by J. Matthews) prepared PDAC tumour models. E. Schmidt produced Siglec-Fc reagents and Z. Jame-Chenarboo performed Siglec-Fc staining on human T cells (advised by M. S. Macauley). M. Kuypers (advised by T. Mallevaey) provided tissues from pet shop mice. D. B. Avila performed ConA and PHA-E/L staining of murine and human T cells. E. S. Y. Chiu expressed and purified recombinant SS. D. Badmaev provided flow cytometry data acquisition support.

H. Cui provided bioinformatics support for bulk RNAseq datasets from murine T cells. L. J. Edgar supervised all aspects of the research. The manuscript was written by L. J. Edgar, F. N. Izzati, H. Choksi, P. Giuliana, and V. Affe with contributions from all other authors.

## RESOURCE AVAILABILITY

### Lead contact

Further information and requests for resources and reagents should directed to and will be fulfilled by the lead contact, Landon J. Edgar.

### Materials availability

This study did not generate new unique reagents.

### Data and code availability

- Bulk RNA-seq dataset has been deposited to the Gene Expression Omnibus (GEO) repository and is publicly available as of the date of publication. Accession number is listed in the key resources table.
- This paper reanalyzes existing, publicly available datasets.
- Code is available upon request
- Any additional information required to reanalyze the data reported in this paper is available from the lead contact upon request.

## EXPERIMENTAL MODEL AND STUDY PARTICIPANT DETAILS

### C57BL/6J Mice

C57BL/6J mice were purchased from Jackson Laboratories or bred at the University of Toronto from Jackson Laboratory founder animals. Mice were maintained under specific-pathogen free (SPF) conditions at the Division of Comparative Medicine, University of Toronto, or at the Biological Sciences Facility, University of Toronto. All animal procedures were approved by the Faculty of Medicine and Pharmacy and Biological Sciences Animal Care Committees (Animal use protocols (AUPs) 20012837 and 20012885, respectively) at the University of Toronto. Unless otherwise noted, 6–12-week-old WT mice were used for all experiments.

### Pet shop mice

Pet shop mice were approximately 7–8-week-old at the time of sacrifice. Mice were purchased from a local supplier High Oaks Ranch and housed in a BioBubble™ containment facility by the Division of Comparative Medicine at the University of Toronto until they were used for experiments. Pet shop mice were purchased under AUP 20012454.

### Pancreatic cancer cell line and culture

CR705 (murine pancreatic cancer) cells^1^ were maintained in RPMI (1.0 g/L glucose), supplemented with 10% v/v heat-inactivated fetal bovine serum (FBS), 1% v/v L-glutamine, and 1% v/v penicillin-streptomycin, (P/S,). Cells were cultured at 37°C, with 100% humidity and 5% CO_2_. When the cells reached a confluency of 80%, they were subcultured. Seventy-two hours before cells were injected into mice, they were seeded at 1 × 10^5^ cells per mL of media in 75 cm^2^ flasks.

### Pancreatic tumor model

Female C57BL/6J (B6; #000664) mice aged 7 to 8 weeks were purchased from the Jackson Laboratory (Farmington, CT, USA). Before mice underwent a single subcutaneous injection of CR705 cells (0.5 x 10^6^ cells in 100 µL *per* mouse), those cells were trypsinized and resuspended in 0.9% sterile saline solution. Each mouse underwent 3-4% isoflurane-based anesthesia before cells were subcutaneously injected into its right flank. From the seventh day onwards, tumors were measured with calipers until they reached a volume of 400 mm^3^ (standard formula π/6 × W2 × L), at which point euthanasia occurred by cervical dislocation, ending the experiment.

### LCMV mouse model

C57BL/6J mice (6-8 weeks old) were purchased from the Princess Margaret Cancer Center or The Jackson Laboratory and handled in accordance with procedures approved by the OCI Animal Care Committee at the Princess Margaret Cancer Center / University Health Network. All mice were housed under specific pathogen– free conditions. LCMV-transgenic mice have been described previously.^2,3^ LCMV-GP_61-80_-specific CD4^+^ TCR transgenic (SMARTA) mice and LCMV-GP_33-41_ specific CD8^+^ TCR transgenic (P14) mice were used.

### LCMV infection and T cell adoptive transfer

SMARTA and P14 T cells were isolated using negative selection (StemCell Technologies) from the spleens of transgenic mice. 1000 P14 and 2500 SMARTA cells were co-transferred intravenously into naive mice via the retroorbital sinus. One day later, the mice were infected intravenously with 2×10^6^ plaque forming units (PFU) of LCMV-Armstrong or LCMV-Clone13 via the retroorbital sinus to induce acute or chronic viral infection, respectively. Virus stocks were prepared and viral titers were quantified as described previously.^4^

### Human PBMCs

Cryopreserved peripheral blood mononuclear cells (PBMCs) from 20–30-year-old healthy individuals were purchased from STEMCELL Technologies (Cat. No. 70025.2) or donated in accordance with the human research ethics board (HREB) biomedical panel at the University of Alberta.

## METHOD DETAILS

### C57BL/6J mice: spleen digestion and PBMC collection

For all C57BL/6J mouse studies, mice were euthanized by carbon dioxide followed by cervical dislocation. Unless otherwise noted, mouse spleens were collected and mechanically digested in FACS buffer (HBSS without Ca^2+^ and Mg^2+^, 1% BSA and 2 mM EDTA) and filtered through 100 µm cell strainer. Red blood cells in the cell suspension were lysed using RBC lysis buffer for 3 minutes and washed with FACS buffer twice. Cells were strained through 40 µm cell strainers prior to analysis by flow cytometry and/or use in T cell stimulation assays.

For mouse PBMC isolation, mouse blood was collected via cardiac puncture immediately after euthanasia into EDTA tubes (BD Biosciences, Cat. 367841). Blood was diluted with an equal volume of PBS and overlaid onto room temperature Histopaque-1083 (Sigma, Cat. 1083-1) in a 2:1 ratio of diluted blood:Histopaque. After centrifugation at 400 x g for 30 mins (room temperature), PBMCs located at the interface between plasma and Histopaque were collected and transferred to a clean tube. Mouse PBMC isolation protocol was kindly provided by Sachiko Kanaji and Taisuke Kanaji (Scripps Research Institute, California).

### Pet shop mice: spleen digestion

Mice were euthanized by carbon dioxide followed by cervical dislocation. Mouse spleens were collected and mechanically dissociated through a 40µm filter in MACS buffer (1X PBS with 0.5% BSA and 2 mM EDTA) into a 50mL tube. After centrifugation for 5 minutes at 1,500 rpm at 4°C, spleen cells were resuspended in 1 mL red blood cell (RBC) lysis buffer (Sigma-Aldrich) and incubated for 6 minutes at room temperature. Samples were washed and resuspended in 1mL of MACS buffer.

### LCMV model: spleen digestion

Mouse spleens for LCMV experiments were digested using the same procedure as WT mouse spleens, except RPMI without supplements was used instead of FACS buffer.

### Pancreatic tumour model: tumor and spleen dissociation

After harvesting, tumors and spleens underwent single-cell processing with a Mouse Tumor Dissociation Kit (#130-096-730; Miltenyi Biotec) and a Mouse Spleen Dissociation Kit (#130-095-929; Miltenyi Biotec), respectively. Both ∼1-2 mm tumor pieces and whole spleens were placed in an ice-cold Tissue Storage Solution (#130-100-008) until being transferred into gentleMACS™ C Tubes (130-093-237; Miltenyi Biotec) loaded with the appropriate kit reagents and enzymes. Then, a gentleMACS Octo Dissociator (#130-096-427; Miltenyi Biotec) ensured the complete dissociation of tumor and spleen samples into single cells. Applying SmartStrainers cleaned samples from debris (30 µm, #130-098-458; 70 µm, #130-098-462; and 100 µm, #130-098-463). Erythrocyte lysis was performed with a Red Blood Cell Lysis Solution (# 130-094-183). The manufacturer provides a detailed protocol for each product.

### Flow cytometry

Unless otherwise noted, the staining for flow cytometry was performed in the following order: viability staining, Fc receptor blocking, surface staining at 37°C, surface staining at room temperature, lectin or Siglec-Fc, fixation and permeabilization, intracellular staining. Cells were stained with Zombie NIR viability dye for 10-15 minutes at room temperature (Biolegend, Cat. 423106, 1:1000 dilution), followed by quenching and washing with FACS buffer. For human Siglec-Fc experiments, Propidium Iodide (PI) was used as the viability dye, instead of Zombie NIR. Fc-receptor blocking was performed for 15 minutes on ice (Biolegend, Cat. 422302 (human), 101320 (mouse), 1:250 dilution). Surface antibody staining was performed at room temperature or 37°C for 30 minutes. Lectin staining was performed on ice for 30 minutes. Intracellular staining was performed using the eBioscience™ Foxp3 / Transcription Factor Staining Buffer Set (Invitrogen, Cat. 00-5523-00) at room temperature for 1 hour. Antibodies used and staining conditions are indicated in the resource table. During panel development, new antibodies were first stained at room temperature for 30 minutes at 1 in 500 dilution (v/v). If staining was not optimal, antibody concentration was titrated and staining at 37°C was attempted to identify appropriate staining conditions. Fixation and permeabilization was performed according to manufacturer instructions. Mouse splenocytes were centrifuged at 350 x g for 5 minutes, and mouse and human PBMCs were centrifuged at 500 x g for 10 minutes at 4°C during staining and washing.

Prior to lectin staining, biotinylated lectins were pre-complexed with FITC-streptavidin (1:1 lectin:streptavidin (v/v)) for at least 15 minutes on ice in lectin staining buffer (HBSS with Ca^2+^ and Mg^2+^, 1% BSA). For mouse Siglec-Fc experiments, 300 nM Siglec Fc was precomplexed with 100 nM Goat anti-human IgG secondary antibody (estimated MW 150 kDa), AF647 (Thermo Fisher, Cat. A-21445) for at least 15 minutes on ice in lectin staining buffer. For human Siglec-Fc experiments, Siglec-Fc chimeras were pre-complexed with 200 nM of Strep-Tactin (1:5 ratio of monomer Strep-Tactin: Siglec-Fc) for 30 min on ice.

Flow cytometry analysis was performed on Cytek Aurora (3 lasers, violet (405 nm), blue (488 nm), and red (640 nm)) and BD FACS Diva. Single color controls for antibodies were prepared using UltraComp eBeads (Invitrogen). Cells were used for Cell Trace Violet and Zombie NIR single color controls. Laser detector settings were established automatically on the Cytek Aurora during Quality Control (QC) using Lot. 2005 QC beads (performed daily). Data was analyzed using FlowJo software (V10.10.0, BD Biosciences) with the UMAP plugin^5^ for dimensionality reduction. Cell sorting was performed on BD FACS Aria with assistance from the University of Toronto flow cytometry core facility staff.

### Siglec-Fc chimeras

Siglec-Fc chimeras were prepared as reported previously.^6^ Briefly, Chinese hamster ovary (CHO) cells were stably transfected with Siglec Fc constructs. CHO cells were cultured for 10 days post-confluency and then the media was collected, centrifuged at 300 x g and filtered through a 0.2 μM filter. The Siglec-Fc was then purified using a HisTrap^TM^ Excel column followed by a Strep-Tactin® Superflow® column and dialysed against PBS at 4°C. Following dialysis, the Siglec-Fc was concentrated using a 50 kDa MWCO Amicon Ultra Centrifugal Filter and then aliquoted into 5 μg aliquouts, lyophilized and the dry Siglec-Fcs were stored at -20°C.

### 3FaxNeu5Ac and DANA synthesis and purification

Methyl (5-acetamido-4,7,8,9-tetra-O-acetyl-3,5-dideoxy-2,6-anhydro-D-glycero-D-galacto-non-2-enopyranos)onate was synthesized according to the previously reported procedure.^7^ This intermediate was then de-esterified and de-acetylated under Zemplén conditions. 3F_ax_Neu5Ac was prepared according to the method reported by Burkart et al.^8^

### T cell stimulation

Cell suspensions (mouse splenocytes or human PBMCs) were prepared at a concentration of 10 million cells/mL in PBS and stained with Cell Trace Violet (CTV) (1:1500 dilution) for 10 minutes in the dark. Pre-warmed cell culture media was added to quench the staining. Cells were cultured in a 96-well round bottom plate at a density of 50,000-600,000 cells/300 µL with 100 U/mL IL-2 (Biolegend, Cat. 575404 (mouse) 589104 (human)). Percentage of T cells in the cell suspensions were quantified using flow cytometry and dynabeads were added accordingly (1:1 bead:T cell ratio for mouse, and 3:1 bead:T cell ratio for humans) (Thermo Fisher, Cat. 11131D (Human), 11453D (Mouse)). Cells were stained and analyzed via flow cytometry after 72 hours unless otherwise noted. For metabolic inhibitor assays, 3F_ax_Neu5Ac and DANA were added to the culture at 100 uM. As proliferation of human T cells was variable, only samples with at least two daughter generations were included in analysis. For cytokine/granzyme analysis, cells were analyzed with and without stimulation with PMA/Ionomycin for 6 hours in the presence of 100 U/mL IL-2 and 10 µg/mL Brefeldin-A.

For Human T cell proliferation experiments with Siglec-Fc chimeras, T cells from three donors were isolated using human T cell isolation kit (Miltenyi Biotech, Cat. 130096535). Isolated T cells were resuspended in flow buffer (PBS with 0.1% BSA/EDTA) and stained with 5 μM (working concentration) of CTV for 7 min at 37 °C and in the dark. 5 mL of Pre-warmed PBS was added to dilute the dye, followed by centrifugation and addition of pre-warmed FBS and PBS (1 mL each) to remove unbound dye. 2 mL of pre-warmed RPMI (10% FBS, 100 Units/mL Penicillin, 100 μg/mL Streptomycin, βME) was added to T cells. The number of T cells for each donor were measured and dynabeads were added in a 1:1 bead: T cell ratio (Thermo Fisher, Cat. 11161D). T cells were cultured in a 96-well round bottom plate at the density of 50,000 cells/200 µL with 100 U/mL IL-2 (BioLegend, Cat. 589104) for 96 hours. All experiments were approved by the human research ethics board (HREB) biomedical panel at the University of Alberta.

### Sialidase treatment

*Streptococcus pneumoniae* (SS) sialidase was expressed with a His-tag in *BL21 E. coli* and purified via Nickel NTA (BioRad, Cat. 12009286) Fast Protein Liquid Chromatography (FPLC). Specific enzymatic activity was measured using the 4-MU-NANA assay^9^. Sialidase treatment was performed at 37°C for 45 minutes in HBSS with 20mM MgCl and 3mM HEPES (pH 7.5). Heat Inactivated controls were prepared by denaturing the enzyme for 30 minutes at 60-65°C.

### Bulk RNA-Seq

Splenocytes from four 1 year 1 month old female mice were used for sequencing. 100 SNA^High^ and SNA^Low^ cells per mouse from CD4^+^ and CD8^+^ T_EM_ (CD44^+^ CD62L^−^), respectively, were sorted into a 96-well PCR plate. Cells were immediately lysed with NEBNext cell lysis buffer with murine RNAse inhibitor from the NEBNext Single Cell/Low Input RNA Library prep kit for Illumina (cat no. E6420S) and stored at −80°C until cDNA library preparation and sequencing at the Donnelly Sequencing Centre (University of Toronto). Resulting reads were trimmed with Trimgalore,^10^ aligned to Mm39 with Star,^11^ and differentially expressed genes were identified with DESeq2,^12^ version 1.38.3.

### scRNA-Seq Analyses

CD4^+^ scRNA-Seq data of four murine spleen samples (i.e. acute/chronic activated and non-activated CD4^+^ T cells) was downloaded from Gene Expression Omnibus (GEO, samples accession numbers: GSM5503101, GSM5503102, GSM5503103, GSM5503104). Murine scRNA-Seq data of CD8^+^ T cells (i.e. P14 naïve, acute and chronic CD8^+^ T cells) was downloaded from GEO (GSE131535). Human pbmc scRNA-Seq data was downloaded from https://www.covid19cellatlas.org/index.patient.html as an h5ad file listed under the ‘Human SARS-CoV-2 challenge resolves local and systemic response dynamics’ section of the Teichmann group.

Quality control: all data was imported into Seurat objects of the Seurat package (v5.0.1) in R (4.3.2). To exclude doublets, poor-quality and dead cells from Murine CD4^+^ T cells, we filtered out cells expressing < 200 genes, cells having a z-score of the number of detected genes < -2 and > 2, and cells with mitochondrial percentage > the top 5^th^ percentile of the mitochondrial genes percentage per sample. For CD8^+^ T cells taken from Murine samples, cells with detected genes < or > 8 median-absolute-deviations (mads) away from the sample median, cells expressing < 200 genes and cells with mitochondrial genes percentage > 8 mads away from sample median were filtered out. For both CD4^+^ and CD8^+^ T cells in mouse, cells contaminated with blood (expressing hemoglobin genes), having log10GenesPerUMI < 0.8 or having ribosomal genes percentage < 5% were also filtered out. Genes expressing at least 1 read in less than 3 cells or genes expressed in less than 10 cells in total were filtered out. Human CD4^+^ and CD8^+^ T cells data used in this study have already passed quality control, have been preprocessed, clustered and annotated in Lindeboom et al. (PubmedID: 38898278).^13^

Preprocessing: each of the mouse samples of CD4^+^ and CD8^+^ T cells were log normalized apart using the NormalizeData function, the top 2000 variable genes were then identified using the FindVariableFeatures function. To prevent cell cycle, mitochondrial and ribosomal genes from driving clustering results, those genes were filtered out from the top 2000 variable genes, then genes expression was standardized using the ScaleData function.

Samples integration, cells clustering and cell type annotation: batch effect correction, also called samples integration, was performed for CD4^+^ Mouse samples apart and CD8^+^ Mouse samples apart using the Harmony method as implemented in Seurat performed with default parameters. Cell neighbours were then identified using 40 dimensions. A resolution of 0.1 and 0.7 were then used to cluster CD4^+^ and CD8^+^ T cells, respectively. UMAPs for each dataset were generated using 40 dimensions. Non-T-cells clusters (i.e. clusters 7 and 9 in Mouse CD4^+^ T cells and cluster 9 from Mouse CD8^+^ T cells) were excluded from downstream analyses. Then each dataset was re-clustered with 40 dimensions and a 0.1 resolution. Then cell types of clusters were annotated by examining the top 10 significantly up-regulated markers of each cluster. Finally, differentially expressed glycan related genes (DEGs) were identified using the FindMarkers function from Seurat with logfc.threshold=1.8 and min.pct=-Inf. DEGs comparisons were performed for activated vs naïve cells in each of CD4^+^ and CD8^+^ T cells in Mouse and Human.

## QUANTIFICATION AND STATISTICAL ANALYSIS

### Statistical analysis and reproducibility

Statistical tests were performed using R for RNA sequencing datasets and GraphPad Prism for all other datasets. All experiments were performed with at least 3 biological replicates. MFI values were only analyzed for cell populations with at least 50 cells. Paired analysis was performed as appropriate. Mixed-effects analysis was performed when samples were paired but some matching data points were excluded (e.g., MFI values for cell populations with <50 cells). Specific n values and statistical tests used can be found in figure captions. Significance was defined as: not significant (ns) ≥ 0.05; *P < 0.05; **P < 0.01; ***P < 0.001.

