## Supplemental Figures for "A Unified Atlas of T cell Glycophysiology"

**Table of Contents:**

Figure S1. Antigen exposure influences glycocalyx presentation S2–S3
Figure S2. Siglec and siglec ligand expression in murine immune cells S4
Figure S3. Phenotypic differences of SNA$^{High}$ and SNA$^{Low}$ cells S5
Figure S4. Persistent T cell activation results in dramatic glycocalyx remodelling and PD-1 expression in antigen experienced T cell populations S6
Figure S5. Representative histograms and quantification of *Phaseolus vulgaris* erythroagglutinin (PHA-E), *Phaseolus vulgaris* leucoagglutinin (PHA-L), Concanavalin A (ConA), and *Galanthus nivalis* Lectin (GNL), staining on murine and human T cells S7
Figure S6. Gating strategy and Siglec-Fc/HAA staining profiles of human T cells from within a mixture of PBMCs S8
Figure S7. Neuraminidase from *S. pneumoniae* efficiently removes α2,3-linked sialic acid from the surface of both mouse and human T cells S9

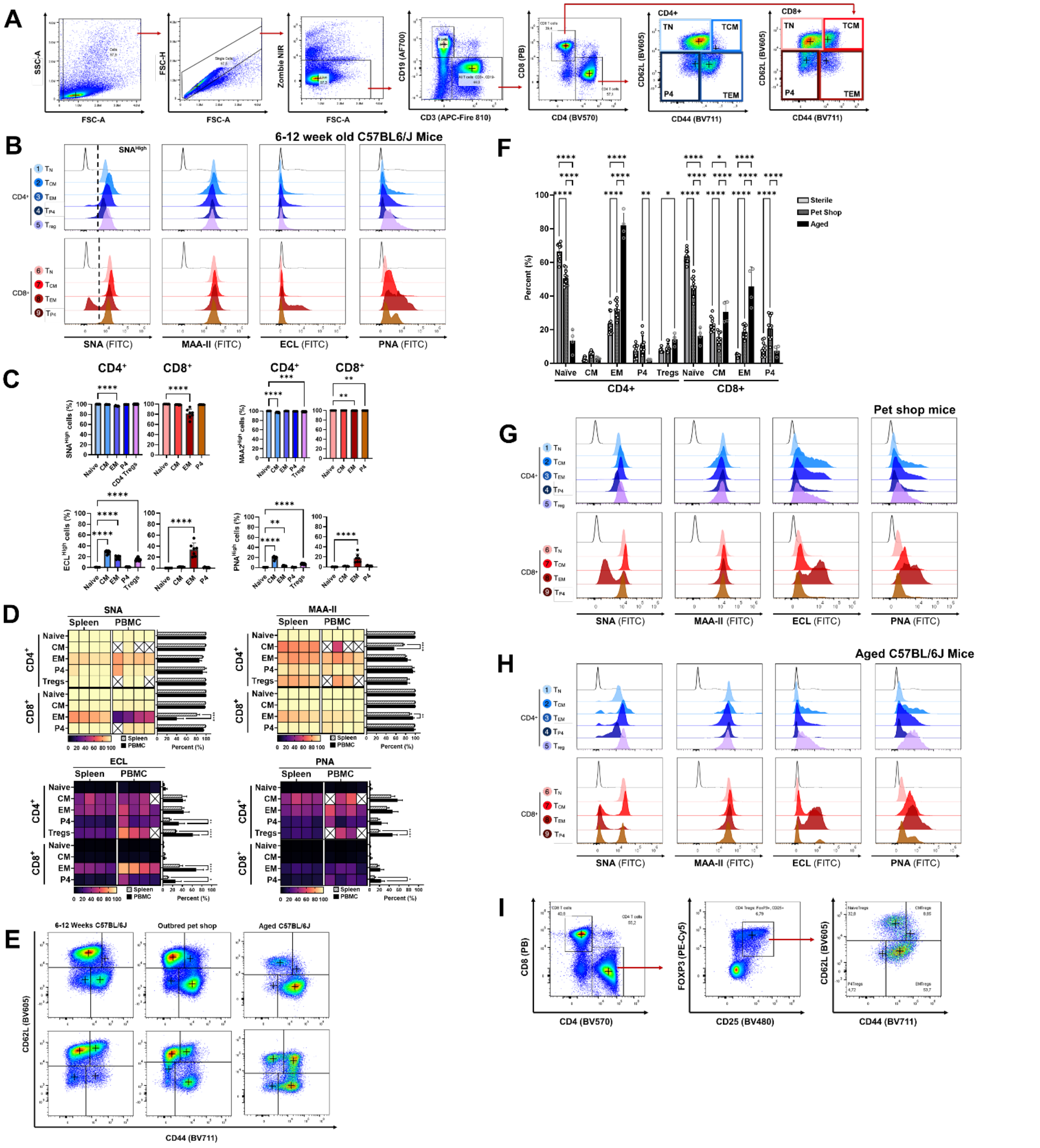


**Figure S1. Antigen exposure influences glycocalyx presentation. (A)** Gating strategy for murine flow cytometry analysis. **(B)** Representative plant lectin histograms from 6–12-week-old C57BL/6J mice (N = 7). **(C)** Quantification of data from **B** presented as Mean ± SD using One-way ANOVA followed by Tukey's multiple comparisons test. **(D)** Heatmap of lectin binding on T cells from matched SPF C57BL/6J mouse splenocytes and PBMCs (percent lectin positive). **(E)** Comparison of CD44 vs CD62L presentation from SPF, pet shop, and aged mice. Data are presented as representative flow cytometry plots. **(F)** Quantification of **E** presented as Mean ± SD using Two-way ANOVA followed by Tukey multiple comparisons test. **(G)** Representative plant lectin histogram from pet shop mice (N = 10). **(H)**

Representative plant lectin histogram from aged C57BL/6J mice (N = 4). Yellow indicates 100% cells positive for lectin staining. Black indicates 0% cells positive for lectin staining. ‘X’ indicates populations with <50 cells collected. Columns represent individual biological replicates (N = 4). **(I)** Gating scheme for $CD4^+$ $T_{reg}$ populations. *$P \leq 0.05$, **$P \leq 0.01$, ***$P \leq 0.001$, ****$P \leq 0.0001$.

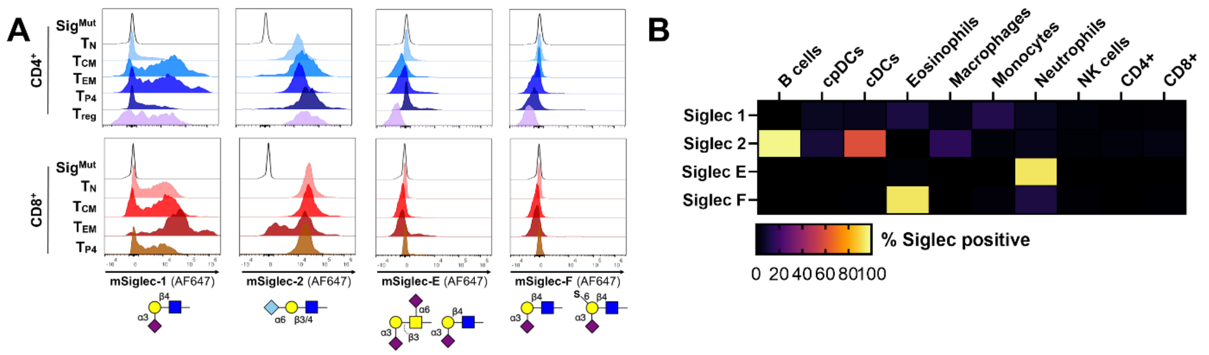


**Figure S2. Siglec and siglec ligand expression in murine immune cells. (A)** Representative histograms of murine T cells stained with mSiglec-Fc reagents (N = 3). **(B)** Heatmap of Siglec expression on murine immune cells. Yellow indicates 100% cells positive for a given Siglec as reported by the anti-Siglec antibodies indicated on the left. Black indicates 0% cells positive for siglec staining. % positive values reported as an average of biological replicates (N = 10)**.**

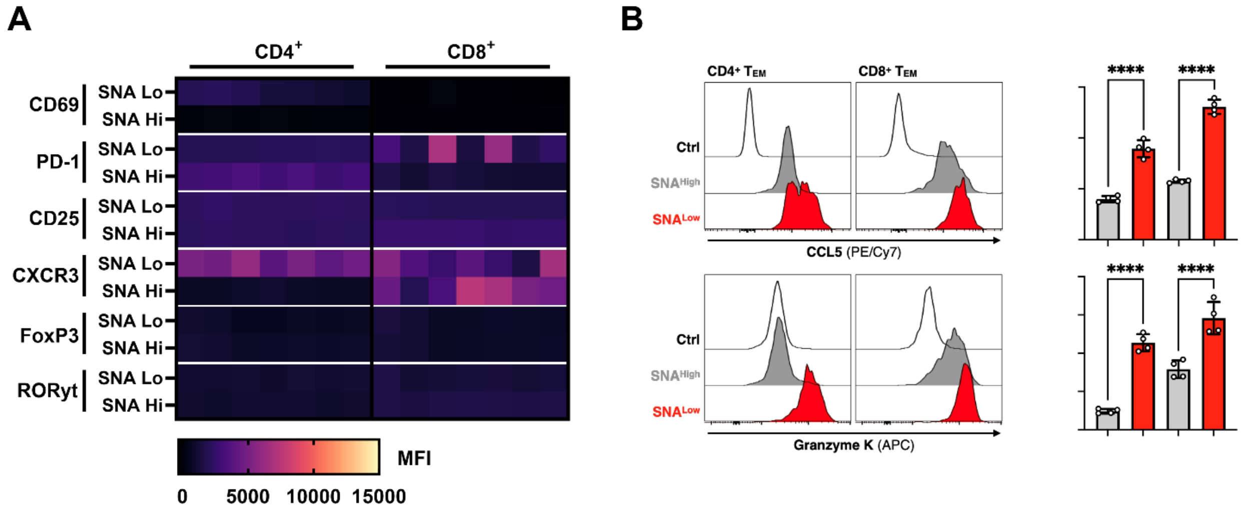


**Figure S3. Phenotypic differences of SNA$^{High}$ and SNA$^{Low}$ cells. (A)** Staining for additional T cell phenotyping proteins in/on SNA$^{High}$ and SNA$^{Low}$ CD4$^{+}$ and CD8$^{+}$ T$_{EM}$ (CD44$^{+}$CD62L$^{-}$) from SPF C57BL/6J spleens (N = 7). **(B)** Representative histograms and quantification of CCL5 and Granzyme K in SNA$^{High}$ and SNA$^{Low}$ CD4$^{+}$ and CD8$^{+}$ T$_{Em}$s from SPF C57BL/6J spleens. Presented as Mean ± SD (N = 4) ****P ≤ 0.0001 using One-way ANOVA followed by Šídák's multiple comparisons test.

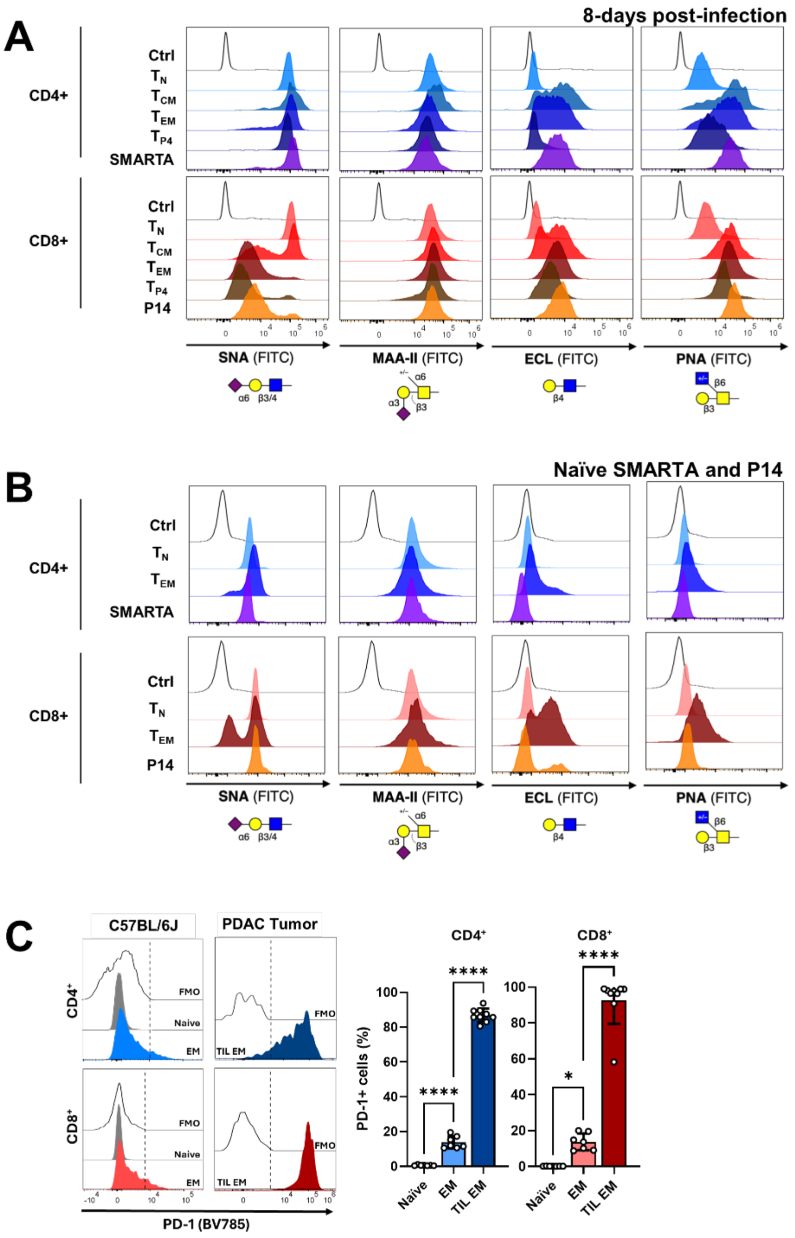


**Figure S4. Persistent T cell activation results in dramatic glycocalyx remodelling and PD-1 expression in antigen experienced T cell populations. (A)** Representative plant lectin histogram from 8 days post-infection with the Armstrong (acute) strain of LCMV (N = 5). **(B)** Naïve SMARTA and P14 shows the same glycosylation pattern with Naïve CD4$^{+}$ and CD8$^{+}$. **(C)** PD-1 expression in C57BL/6J Naïve and Effector Memory T cells (EM) from spleens of untreated mice, and tumour infiltrating lymphocytes effector memory (TIL EM) T cells from PDAC mouse tumors. Presented as Mean ± SD (N = 7 for untreated animals and N = 9 for TILs). *$P \leq 0.05$, **$P \leq 0.01$, ***$P \leq 0.001$, ****$P \leq 0.0001$ using One-way ANOVA followed by Tukey's multiple comparisons test.

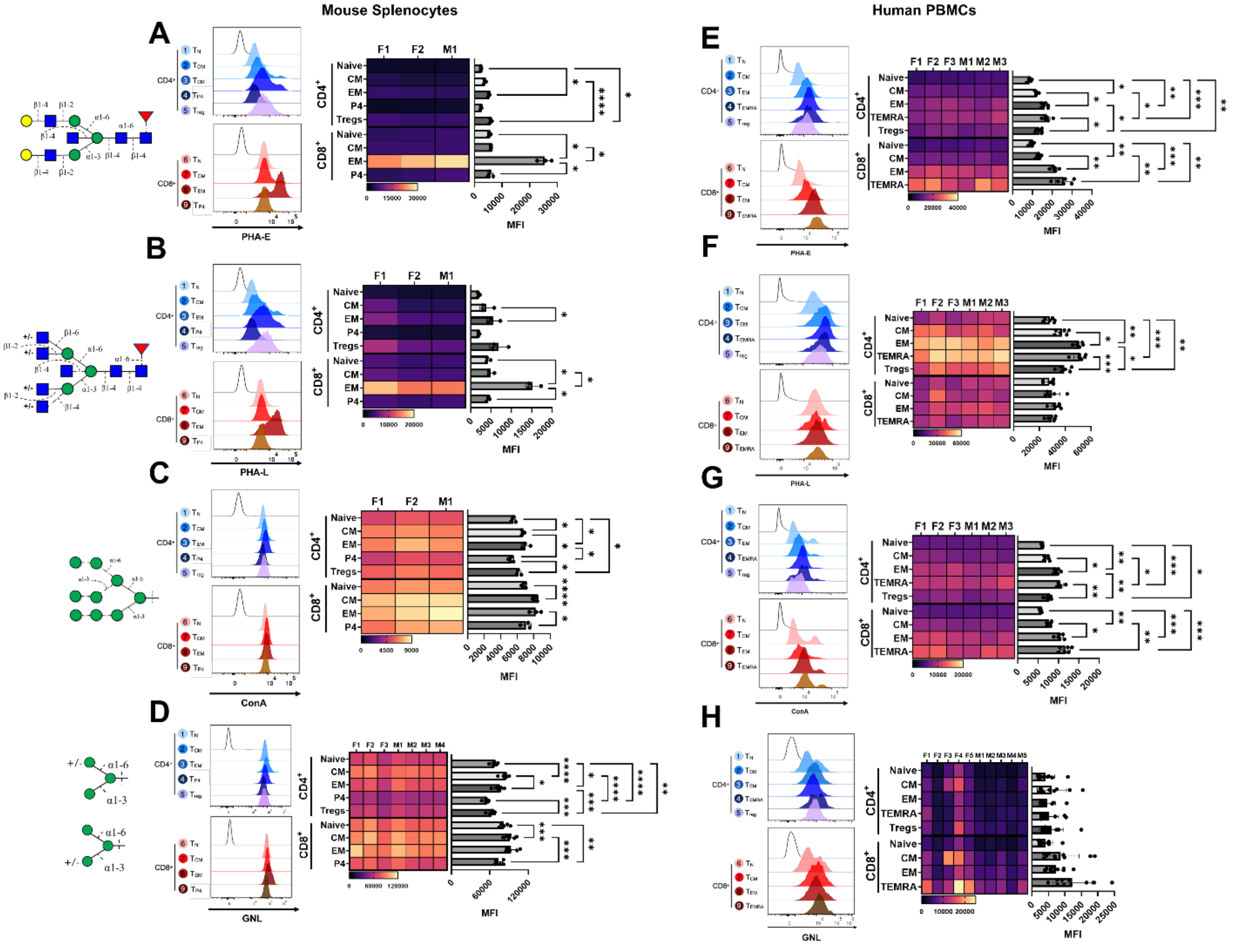


**Figure S5. Representative histograms and quantification of *Phaseolus vulgaris* erythroagglutinin (PHA-E), *Phaseolus vulgaris* leucoagglutinin (PHA-L), Concanavalin A (ConA), and *Galanthus nivalis* Lectin (GNL), staining on murine (A-D) and human (E-H) T cells.** Presented as Mean ± SD (N = 3) *P ≤ 0.05, **P ≤ 0.01, ***P ≤ 0.001, ****P ≤ 0.0001 using One-way ANOVA followed by Tukey's multiple comparisons test.

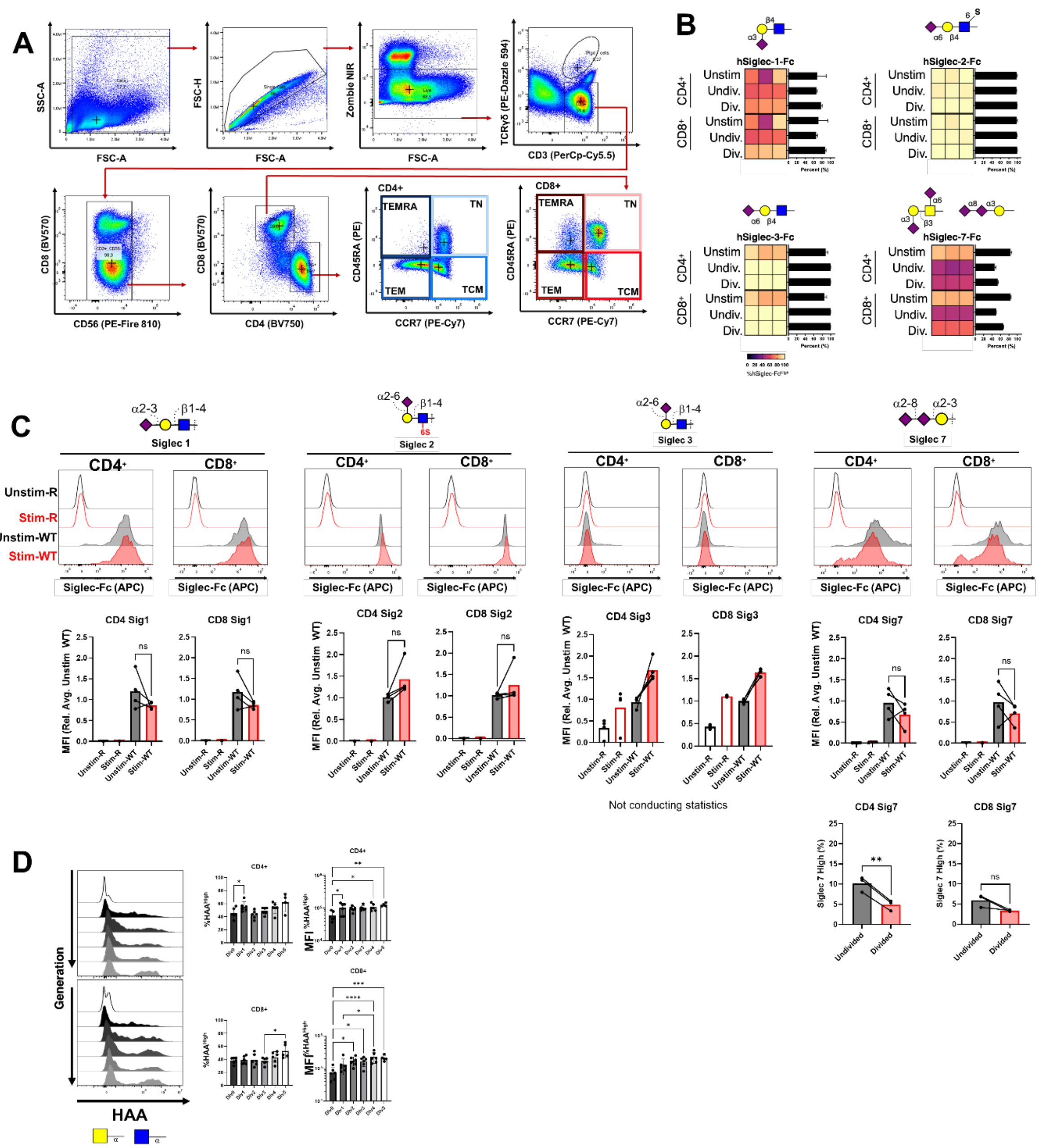


**Figure S6**. **Glycan profile of activated human T cells with Siglec Fc and other lectin detection reagents. (A)** Gating strategy for human PBMC flow cytometry analysis. **(B)** hSiglec-Fc binding on resting **(B)** and proliferating **(C)** $CD4^+$ and $CD8^+$ human T cells. '-R' denotes staining control, '-WT' denotes WT hSiglec Fc. **(D)** HAA lectin binding on proliferating $CD4^+$ and $CD8^+$ human T cells. Presented as Mean ± SD (N=10). *P ≤ 0.05 using mixed-effect analysis. *P ≤ 0.05, **P ≤ 0.01, ***P ≤ 0.001

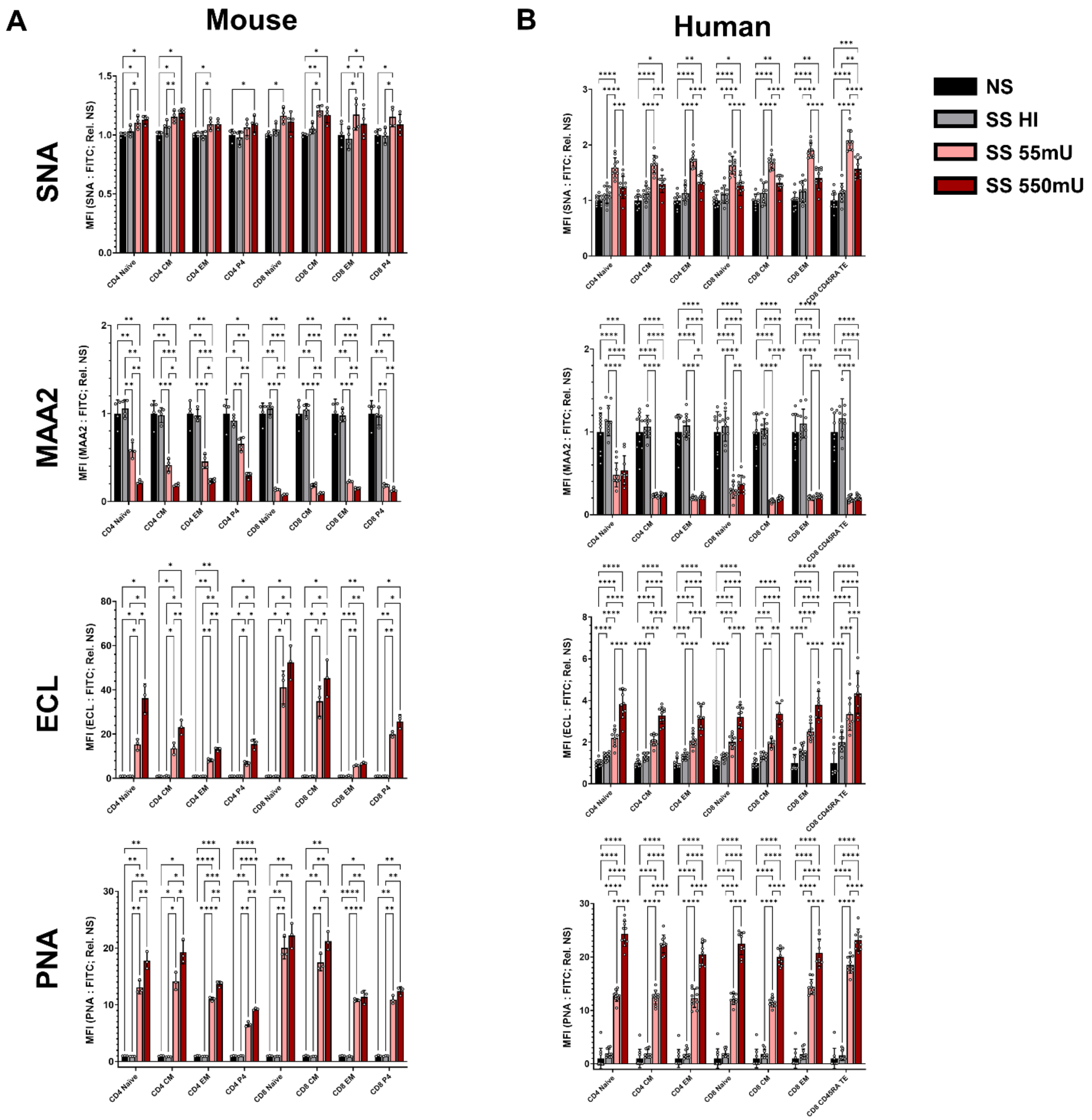


**Figure S7. Neuraminidase from *S. pneumoniae* efficiently removes α2,3-linked sialic acid from the surface of both mouse (A) and human (B) T cells.** Presented as Mean ± SD (N=10) *P ≤ 0.05, **P ≤ 0.01, ***P ≤ 0.001, ****P ≤ 0.0001 using Two-way ANOVA followed by Šídák's multiple comparisons test.
